# The NMR Exchange Format (NEF): Specification and Applications

**DOI:** 10.64898/2026.04.22.715536

**Authors:** Eliza Płoskoń, Kumaran Baskaran, Roberto Tejero, Charles D. Schwieters, Benjamin Bardiaux, Peter Guentert, Rasmus H. Fogh, Aleksandras Gutmanas, Edward J. Brooksbank, Masashi Yokochi, David S Wishart, Jonathan R Wedell, Wim F. Vranken, Daniel Thompson, Gary Thomson, Brian O. Smith, Saima Rehman, Theresa A. Ramelot, Timothy J. Ragan, Alberto Perez, Binod L. Perera, Ezra Peisach, Michael Nilges, Luca G. Mureddu, Arup Mondal, Emilia A. Lubecka, Adam Liwo, Genji Kurisu, Naohiro Kobayashi, Piotr Klukowski, Bruce A. Johnson, Yuanpeng J. Huang, Jeffrey C. Hoch, Victoria A. Higman, Torsten Herrmann, Morgan W. Hayward, James A. Garnett, David A. Case, Stephen K. Burley, Paul D Adams, Gaetano T. Montelione, Geerten W. Vuister

## Abstract

The NMR Exchange Format (NEF) is a community-driven standard for representing NMR experimental data in a consistent, interoperable, and machine-readable form. Built on the STAR syntax, NEF provides a structured framework for storing and exchanging chemical shifts, peak lists, various types of structural restraints, and related metadata, thus allowing for data exchange across software platforms. By enabling direct, lossless transfer of information, NEF simplifies multi-software workflows, improves reproducibility, and supports FAIR (Findable, Accessible, Interoperable, Reusable) data principles. We describe the NEF specification, its current implementation across commonly used NMR software packages, and its application in areas including biomolecular structure determination, metabolomics, and ligand screening. Testing demonstrates that NEF can be used to exchange complete datasets between programs without loss of information or functionality. We also outline recent developments and future directions, such as inclusion of NMR relaxation data and support for non-standard residue topologies. NEFs growing adoption highlights its potential as a unifying standard for NMR data, enabling more efficient, transparent and collaborative research.

## Introduction

Nuclear Magnetic Resonance (NMR) methodologies allow for the study of molecular properties including structure, interactions, assembly and dynamics. Historically, the parallel development of dedicated software for analysing different kinds of NMR data has resulted in a proliferation of software-specific and incompatible file formats, which limits scientific data interoperability and reuse. Previous attempts towards unifying NMR data formats, like NMR-STAR (*1*) or the CcpNmr data model (*2*), partially succeeded to achieve data interoperability but failed to gain general community uptake. Addressing the challenge of NMR data interoperability crucially depends on the development and adoption of a lightweight file format that adheres to the FAIR principles (Findable, Accessible, Interoperable, and Reusable) while maintaining the flexibility required to accommodate diverse data types and future expansion (*3*). To address the various challenges facing NMR data analysis, the NMR community has developed the NMR Exchange Format (NEF) through a series of collaborative meetings and development working groups (*4*).

A unified, FAIR-compliant file format alleviates the need for data converters between various formats. Despite several attempts to develop these, data loss proved to be an inherent challenge (*2, 5*). Several documented cases have shown that critical details can be lost during such format conversions(*6*), significantly affecting the accuracy of scientific analysis and leading to potential misinterpretations. For example, in absence of universally defined ontologies for NMR-related data, even fundamental parameters such as ‘peak intensity’ can be defined in different ways, e.g. it can be quantified either as peak-integral or peak-height. These different definitions can substantially influence data analysis and calculations, highlighting the crucial need for consistent and clearly interpretable definitions of such quantities. The lack of standardisation also complicates, and sometimes prevents, validation between models and experimental data, and contributes to the perception of NMR-derived molecular structures as being inferior to those derived from X-ray crystallography (*7*).

The NEF development is an ongoing effort designed to address these problems based on two essential principles: a minimal, unambiguously defined mandatory core set of information (*vide infra*) that all NEF-compatible programs must support, and a flexible, extensible architecture that accommodates new and emerging data types and software-specific information. Importantly, the latter information can be ignored by programs without changing the file’s usability or readability for other software packages, while still being available if needed. The pass-through of all data is encouraged but not mandated.

This paper documents NEF’s current technical specifications and demonstrates its practical implementation across multiple software platforms. We present detailed format specifications, validate NEF’s effectiveness through comparative benchmarking of major structure calculation programs, and demonstrate its integration into current NMR workflows. We also examine NEF’s growing adoption within the community and discuss ongoing developments that extend its utility beyond traditional structure determination applications.

## Results

### NEF file format and syntax

NEF is based on the STAR (Self-defining Text Archive and Retrieval (*8, 9*)), which is well-established in crystallography and NMR. This syntax is also used in PDBx/mmCIF files, the preferred archival data format for the world-wide Protein Data Bank (*10*) (wwPDB) atomic coordinate data (*11*), and in NMR-STAR files, which are the archive format for NMR data in the Biological Magnetic Resonance Bank (BMRB) (*1*). An example NEF file is shown in Fig. 1; a fully annotated NEF example file is available on the NEF GitHub site (https://github.com/NMRExchangeFormat/NEF/).

**Figure 1.**
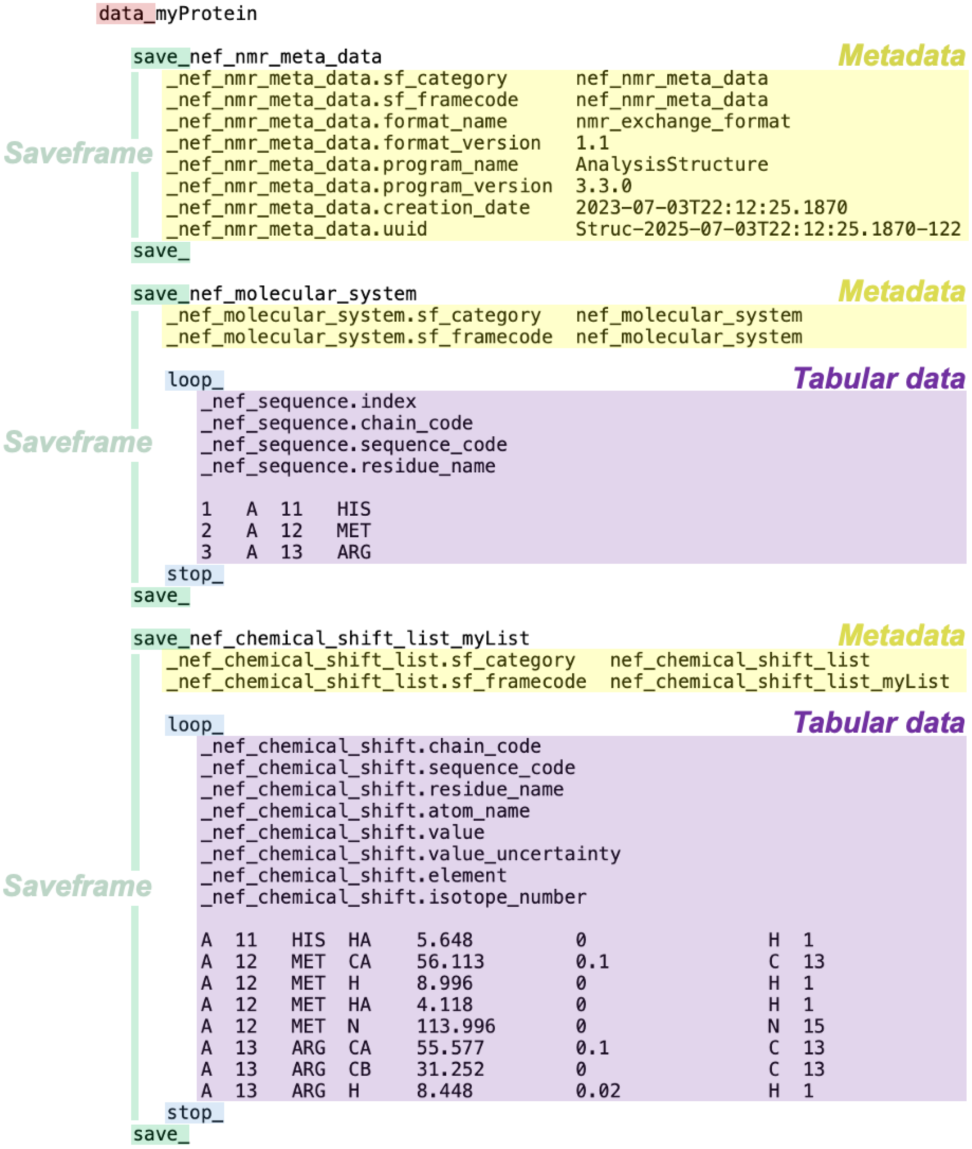
Overview of a NEF file. The example illustrates three saveframes types: `_nef_nmr_meta_data` (top), `_nef_molecular_system` (middle), and `_nef_chemical_shift_list` (bottom); note that the chemical shift list in this example is named ‘myList’. Metadata blocks (yellow) define general information relevant to each saveframe. Tabular data blocks (purple) contain structured data entries, including molecular system components and chemical shift assignments. Note that the latter two saveframes will generally be much more extensive than the example shown.

Like all STAR format files, NEF files are self-describing. Each data element is paired with a tag (data name), forming a tag-value pair. Tags are defined in an associated, but separate, data dictionary (https://mmcif.wwpdb.org/dictionaries/mmcif_nef.dic/Index/) which specifies the format of the expected values as well as their mandatory or optional inclusion status. Notably, the NEF dictionary itself is defined using a STAR syntax as well.

Tags in NEF have a specific namespace, category and tag-name structure consisting of two parts separated by a full stop. The first part starts with an underscore (‘_’), making them uniquely identifiable within the file structure. It identifies the saveframe or loop category. Tags specific to NEF-defined parameters begin with `_nef`, e.g. `_nef_nmr_spectrum`, effectively defining a NEF-specific namespace. Custom namespaces can be used by other programs (*vide infra*). The second part of the tag denotes the tagname and provides additional information, e.g. the spectrum experiment type as in `_nef_nmr_spectrum.experiment_type`.

NEF files are text-based and use the extended STAR character set to store data elements separated by whitespace. Line breaks are allowed for multiline strings. NEF files use four reserved keywords: `data_`, `loop_`, `save_`, and `stop_`. The file starts with the `data_` keyword followed by the project name, initiating the single data block contained within the file. Each NEF file must contain exactly one data block.

The single data block of the NEF file is organised into sections called saveframes, which allows for a clear separation of the storage of data for the various elements used for NMR data analysis e.g. peak lists, shift lists, restraints lists, etc. Each saveframe starts with the `save_` keyword prefix concatenated to a saveframe identifier consisting of a namespace followed by a category and name. For example, `save_nef_chemical_shift_list_myList`, defines a chemical shift list in the NEF namespace with the name “myList”. Saveframe names must be unique within a NEF file. The end of the data contained in a saveframe is indicated by a standalone `save_` keyword. The saveframe contains a mandatory metadata section with at minimum two tags with tag names `sf_category` and `sf_framecode,` prefixed by the type of the saveframe category. These two tags together serve to identify the category and name of a specific saveframe. Unique singleton saveframes, such as the molecular system or metadata, do not have a name and for these the values of `sf_framecode` and `sf_category` tags are identical. Additional saveframe-specific mandatory and non-mandatory metadata tags provide contextual information about its content. For instance, the metadata of the peak list saveframe (*vide infra*) must specify the number of peak dimensions and has to contain a reference to the associated chemical shift list, by which peaks can be assigned.

Following the STAR convention, NEF does not allow multiple occurrences of the same tag within the same saveframe. Therefore, to handle lists or tabular data that are a common occurrence in NMR, NEF uses loops. A loop is a collection of tags with multiple values, structured in a space-delimited format, initiated by the `loop_` keyword and terminated by the `stop_` keyword. The tags immediately following the `loop_` delimiter effectively serve as column headers. As with saveframe tags, loop tags specific to NEF begin with the _`nef` namespace and are structured, having both a category and a data name part separated by a full-stop.

NEF allows for booleans, integers, floats, and single-word values, whereas data that include spaces, tabs, or newlines must be enclosed in single (‘) or double (“) quotes (see also Supplementary material “Format-specific details” section). This setup allows for Python-style encoding of quoted text without the need for complicated escape characters. Missing or null data is indicated by a full-stop (“.”).

The NEF specification describes several saveframes for the most common forms of NMR data. These include chemical composition, chemical shift data, peak lists derived from one- or multidimensional experiments, and restraints related to distance, torsion angles and residual dipolar couplings (RDCs). In addition, NEF has a dedicated metadata section that specifies the format’s version and enables data tracking. Depending on the nature of the data it contains, some types of saveframes are restricted to occurring only once per file, e.g. the composition of the molecular system, while others may occur multiple times, e.g. peak lists or chemical shift lists. NEF file requires three mandatory saveframes: `_nef_nmr_meta_data`,`_nef_molecular_system`, and `_nef_chemical_shift_list`. Although the chemical shift list may be empty and still satisfy the format requirements, such a file does not meet the criteria for deposition. We will discuss the key features of various saveframes below.

### Metadata

To ensure data traceability, it is essential to include information that identifies the data source and specifies the versions of both the software and the standard used. NEF files encode this information in a single `_nef_nmr_meta_data` saveframe. This saveframe records key details such as the names and versions of software that created the file or modified it, the NEF file version, the creation date and the file’s universally unique identifier (UUID, see also Supplementary material “Format-specific details” section). The metadata saveframe is designed to also be able to document the entire data management history, with optional loops to store its history or the software tools and scripts used during processing. Additionally, the metadata saveframe supports the inclusion of references to related database entries, thus supporting integrated structural biology approaches in line with the federated model of data management (*12*).

### Sequences, residues and atoms

The molecular composition of the system being studied is recorded within the `save_nef_molecular_system` saveframe using a `_nef_sequence` loop. This type of saveframe only occurs once per NEF file. Residues are identified by three non line-breaking strings (`chain_code`, `sequence_code`, `residue_name`), which must be applied consistently throughout the various saveframes included in the file. In line with the mmCIF format for atomic structural data, NEF uses standard IUPAC case sensitive, uppercase residue names for amino acids and oligonucleotides residues. Sequence codes are strings but cannot contain wildcards, as the NEF standard defines explicit residue identities. For polymer entries, e.g. protein, DNA or RNA, sequence codes are expected to be interpretable as monotonically increasing integers.

The NEF specification includes a list of atom and atom group names for all standard residues derived from IUPAC-defined chemical compound descriptions (*13*), as well as the allowed wildcards (see below for detail). Atoms are identified by adding a fourth case-sensitive string, e.g. ‘HA’, to the residue identifier to yield the four-string (`chain_code`, `sequence_code`, `residue_name, atom_name`) atom identifier. Notably, residue names and atom names that do not match the molecular sequence are allowed in the peak list and chemical shift saveframes (discussed below), as they may refer to observed but as yet unassigned resonances, i.e. actual signals in the NMR experiments.

The residues’ default states are defined by NEF to represent common NMR conditions at pH 7, which assumes protonated lysine and arginine residues and de-protonated aspartic and glutamic acids. Histidine was chosen as neutral, protonated at the N^δ1^ (ND1) position. Residue variants, such as those representing different protonation states or covalent modification, can be annotated using a separate tag (column) in the `_nef_sequence` loop which lists atoms that are either added or omitted (see also Supplementary material “Format-specific details” section). This approach avoids the need for a predefined list of residue variants, which would require constant maintenance, while still enabling accurate representation of residues’ atom composition. Sequential linkages are handled by a separate optional tag (column) distinguishing between ‘start’, ‘middle’, ‘end’, ‘cyclic’, ‘single’ and ‘break’, variants, with the default being a single start-middle-end sequence. Additional covalent linkages, including disulfide bonds, are recorded in the dedicated `_nef_covalent_links` loop, enabling precise specification of chemical bonds between atoms. Finally, a specific boolean `cis_peptide` value is also provided for proteins.

### Chemical shift lists

In NEF, resonance frequencies, i.e. the signals observed in NMR experiments or predicted from computation, are stored in a chemical shift list saveframe. Following the convention of the molecular system saveframe, atoms are identified by the four-string atom identifiers. Although element and isotope codes can *generally* be inferred from the atom name, they are explicitly specified by separate mandatory tags contained in the chemical shift saveframe to avoid ambiguity.

Most resonances can be directly mapped to specific atoms in a structure and can consequently be clearly annotated with IUPAC atom names as also found in structure models. Some prochiral atoms, such as the methylene (CH_2_) hydrogens around a prochiral centre or prochiral methyl groups in leucine or valine residues, cannot be distinguished *a-priori* without specific experiments. In these cases, the assignment must reflect this ambiguity to avoid incorrect labelling, which could affect both structure calculation and structure validation routines, as well as overall functional interpretation.

To address this ambiguity issue, NEF introduces a simple naming system. Atoms with stereospecific assignments use their standard IUPAC names. Non-degenerate, non-stereospecific assignments use the characters ‘x’ and ‘y’, while accidentally degenerate resonances or NMR-equivalent atoms that cannot be distinguished are labelled with the ‘%’ symbol. For example, the two arginine H^β^ atoms might be named HB2 and HB3 (stereospecific and non-degenerate), HBx and HBy (non-stereospecific and non-degenerate), or HB% (equivalent or degenerate).

In contrast to the wildcards (%), NEF avoids expanding pseudoatom names, e.g., ALA MB, into individual atoms, i.e. if used in distance they are to be interpreted to denote the geometric mean of the constituting atoms. The NEF representation for an expanded group would use the wildcard nomenclature, i.e. ALA HB% for the alanine example. Pseudoatoms and wildcards may appear in the same file, yet it is recommended to use the wildcard notation as it implies the physically more relevant interpretation.

The use of non-unique atom designators, rather than standard IUPAC ones, allows conversion to the related NMR-STAR format but not the reverse.^11^ To support transparent mapping of NEF atom names to the IUPAC nomenclature used by atom structural information containing PDBx/mmCIF files, the PDBx/mmCIF dictionary includes a specific tag (`pdbx_atom_ambiguity)` in the _`atom_site` loop to document the corresponding NEF atom name (See Restraints section below).^10^ Ambiguous atom mappings can be explicitly included in the mmCIF coordinate file using this tag when depositing or associating NMR data in NEF format. Note that the legacy PDB format is not able to support these mappings and therefore its usage is not recommended.

### Spectra and Peak Lists

NEF does not store processed spectral data, which inevitably will exist in various binary formats. Efforts to define standardisation of spectral data are currently being initiated by the BMRB. Instead, it supports the storage of peak lists, as peaks provide a representation of the NMR data that underpin nearly all downstream NMR data analysis steps, such as assignments and structure calculations. While essential for such analyses, peak lists have thus-far often been omitted from data depositions due to missing contextual details, like observed isotopes, axis units, or magnetization transfer types connecting the various dimensions. NEF addresses this issue by providing a structured format that requires basic spectral descriptions, ensuring that all the metadata linking peaks to the relevant experiment and chemical shift data are included.

Peaks are identified manually or automatically by software packages and are stored as spectrum-specific peak lists in a named _`nef_nmr_spectrum` saveframe (Supplementary Fig. S1). The saveframe uses the mandatory metadata tag `num_dimensions` to specify the number of spectral dimensions, supported by two required loops: the `_nef_spectrum_dimension` loop describing each observed dimension and the `_nef_spectrum_dimension_transfer` loop defining the magnetization transfer between dimensions. Spectrum descriptions in the dimension loop define the mandatory identifier (id, *unique within the spectrum*) and units for each spectrum dimension, and can include optional details such as the spectral width, the number of points, reference frequencies and whether the dimension is folded. For the dimension-transfer loop, transfer types are selected from seven predefined values, e.g. `through-bond, through-space,` etc., as predefined in the NEF specification. Although in rare cases this model may not fully capture complex magnetization pathways, it offers a broadly applicable and extensible framework for describing NMR experiments. Crucially, NEF avoids dependence on fixed experiment names, thereby sidestepping the need for a centrally maintained and controlled vocabulary, which would be both impractical and unsustainable.

The `_nef_peak` loop in the _`nef_nmr_spectrum` saveframe encodes the actual peak positions, height and/or volumes and their uncertainties, as well as optional assignments. If multiple peak lists relate to the same spectrum, they are stored in separate saveframes and retain distinct names. Since data traceability is a core principle of NEF, each peak is assigned a unique integer `peak_id` within its saveframe. In response to software developers’ requests, these peak identifiers (`peak_id` and `_nef_nmr_spectrum`) should remain consistent across input and output NEF files, enabling downstream data, such as restraints, to be unambiguously linked back to their originating peaks. The linking mechanism is detailed in the “Restraint Links” section (see below). Alongside peak positions, peak intensity is widely used in data analysis. Since “intensity” may refer to either height or volume, each with different implications, NEF distinguishes between these by assigning separate tags for volume and height in its peak definitions.

A _nef_nmr_spectrum saveframe can describe peaks in spectra of any dimensionality up to an arbitrary practical limit of fifteen, as adding a spectral dimension simply requires the addition of an extra column in the _nef_peak loop and an extra line in both the dimension and dimension-transfer loops.

### Restraints: distance, dihedral and RDC

Restraints are essential for defining spatial relationships between atoms and play a critical role in molecular structure determination. Derived from experimental data, they impose limits on distances, dihedral angles or orientations within a three-dimensional model. NEF organizes restraint data into type-specific saveframes for distances, dihedral angles, and RDCs. This structure provides a flexible framework that can be easily extended to support new restraint types from emerging experimental techniques. Restraints are typically generated either directly from experimental data, or by structure refinement programs that use the peak lists and the chemical shifts in iterative cycles of data interpretation and structure calculation.

Each _`nef_restraint_list` saveframe includes a mandatory tag indicating the form of the associated pseudo-energy function used during structure calculation, such as square-well-parabolic, log-normal etc. Additional metadata, such as tensor parameters, must be included depending on the restraint type.

Restraints in NEF are organized in loops that specify the atoms involved, target values, uncertainties, bounds, and weighting. Each restraint is assigned a unique integer `restraint_id` to ensure traceability throughout the structure determination process. Ambiguous restraints, common in NMR when multiple assignments are possible, are represented by multiple rows sharing the same `restraint_id`, *i.e.* the “or” operation, and optionally linked by the tag `restraint_combination_id` which assigns a unique integer defining the “and” operation in a similar fashion. This setup supports multiple alternative interpretation use cases, for example in supporting dihedral restraints with two allowed φ, ψ regions in the Ramachandran dihedral space, while preserving simplicity and data integrity.

A persistent obstacle in depositing restraints and their associated structures has been the handling of non-stereospecific assignments, which varies across different NMR structure calculation programs. For example, to satisfy NOESY data ARIA (*14, 15*) permits floating chirality for stereospecific groups within individual conformers of an ensemble, whereas CYANA (*16, 17*) and Xplor-NIH (*18, 19*) enforce identical assignments across all members of the final ensemble. As a result, using ARIA the methylene protons, or valine and leucine methyl groups, and certain aromatic ring-proton pairs may be dynamically flipped during structure calculation, such that different models within an ensemble can adopt different conformer stereochemistry. Other structure calculation programs also require a mechanism to record their stereochemical mappings.

To implement a general and flexible system for dealing with NMR-related stereochemistry issues in the context of atomic structural information, NEF is compatible with an extension of the PDBx/mmCIF dictionary, by employing the non-mandatory mmCIF tag _`atom_site.pdbx_atom_ambiguity`. Data defined by this tag, together with `_atom_site.auth_seq_id`, `_atom_site.auth_comp_id`, `_atom_site.auth_asym_id`, encodes the atom name identifier, i.e. chain, residue name, residue id and NEF atom name, thus providing a direct mapping between the NEF atom name identifier as used in defining the restraints and the corresponding atomic coordinates (Supplementary Fig. S2). Stereochemistry can thus be unambiguously mapped, e.g. HBx onto HB2 and HBy onto HB3 or *vice versa,* on a per model basis. Adoption of this tag enables unambiguous downstream validation and reproducibility. It also solves a long-standing issue related to the nomenclature of tyrosine and phenylalanine aromatic ring atoms, which is dependent upon their χ_2_ dihedral angle resulting in swapping the CD1/HD1/CE1/HE1 and CD2/HD2/CE2/HE2 atoms when χ_2_ exceeds the ±90° boundaries. Consequently, this may result in major restraint validation errors across conformers in the NMR ensemble.

### Restraint links

Cross-referencing between saveframes is a useful way to connect data elements, but also increases complexity, thus complicating the handling of NEF by software packages. However, it was deemed crucial to maintain the linkage between peaks and restraints, as it subsequently enables powerful data analysis (see examples below). NEF allows for linking of peaks and restraints in a single non-mandatory _`nef_restraint_links` saveframe. Each row in the accompanying loop contains four mandatory tags: the `nmr_spectrum_id` identifying its peaklist saveframe, the `peak_id` from this saveframe, and the corresponding `restraint_list_id` and `restraint_id` from the restraints saveframe. This provides an unambiguous cross-reference between experimental observations and the restraints derived from these, ensuring that within the NEF file all derived data, i.e. a restraint, can be traced back to its source, i.e. a peak or peaks.

### Software-specific NEF extensions

Different software packages may use program-specific parameters or data that are essential for reproducing analyses, which may not be directly usable or relevant for other tools. NEF supports this requirement by allowing extra namespaces to be defined and used in metadata tags, additional loop columns, new loops, or even entirely new saveframes. This ensures compatibility, making these clear for other tools to identify, ignore or use if needed. For example, ARIA 2.3.3 (or higher) uses the ‘_`aria’` namespace for additional data (Supplementary Fig. S3), and CcpNmr Analysis V3 can encode its entire project data within a NEF file using the ‘_`ccpn’` namespace, without affecting the ability of other NEF-compliant software to read the standard NEF content. It is highly recommended that all data supported by NEF is reported using the appropriate NEF tags and that non-NEF namespaces are only used when there is no support for the information provided by NEF. Software developers can request a specific namespace to be reserved for their software; these are documented as part of the NEF specification on the NEF Github website (<u>NEF/specification at master · NMRExchangeFormat/NEF · GitHub</u>) in the form of a commented example and dictionary file. Table 1 lists the namespaces already allocated for the fifteen NEF-supporting software programs reported in this paper. During deposition to the wwPDB only data described by the NEF dictionary, i.e. those without any additional software-specific tags, is included. This ensures the correct annotation of the data within databases and maintains consistency and compatibility across different software platforms.

**Table 1.**
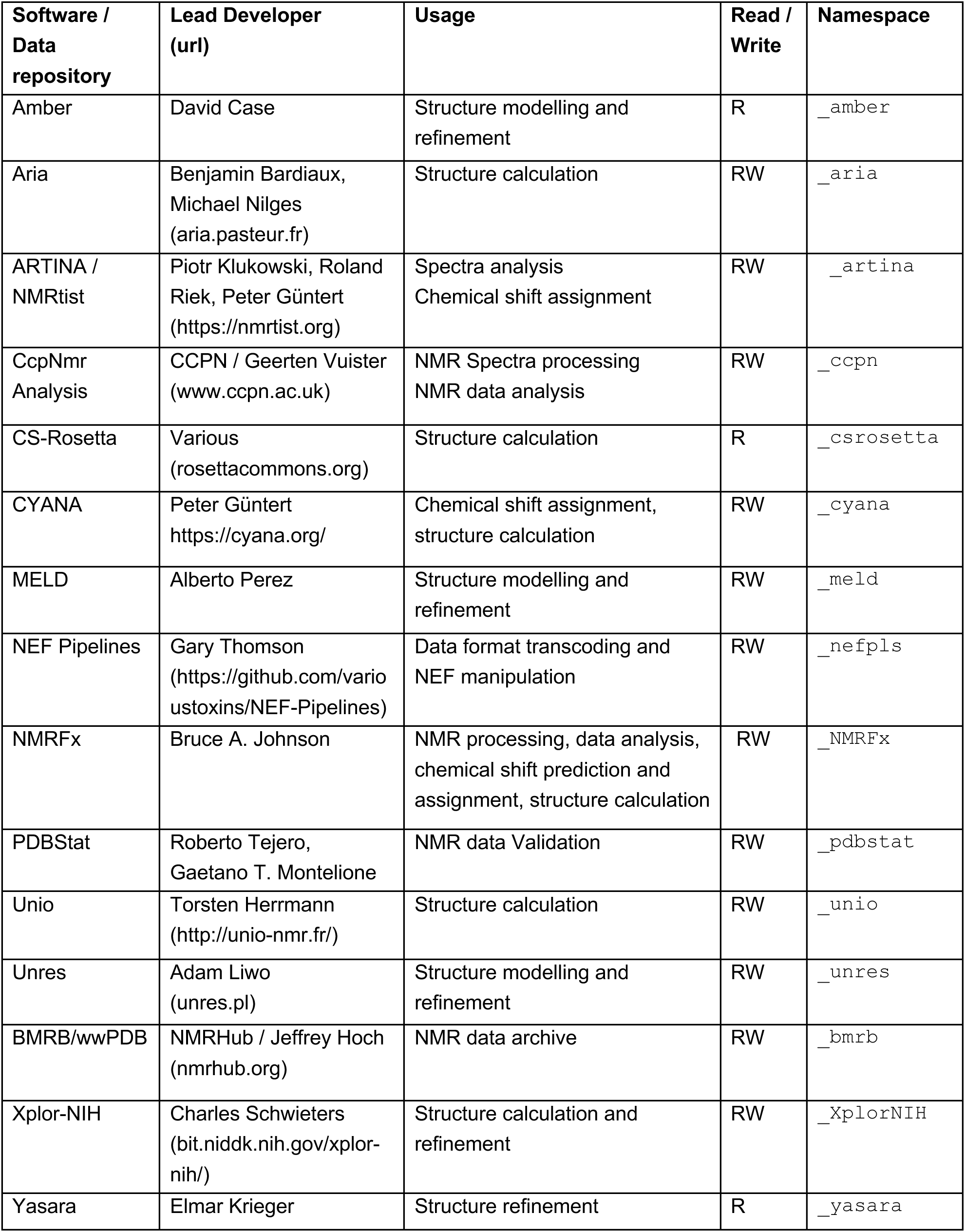
Current adoption of NEF format.

### Testing NEF

The first step in the broader adoption of any new file format is robust testing, focusing on the core data reading and writing functionalities without loss of information. Interoperability testing follows the best practices established in software engineering and can help identify errors before they impact end users. To demonstrate NEF’s applicability across an extended NMR software ecosystem, we processed NEF datasets of fifteen representative proteins (Supplementary Table ST1) with each of the various software programs (Table 1). These test cases were selected to cover both common and more challenging use cases across a range of molecular sizes, data content and restraint complexities, thus providing a varied set of examples to evaluate software compatibility representative of a cross-section of scenarios commonly encountered in biomolecular NMR workflows with an initial emphasis on protein calculation as a starting point.

The testing sets span protein lengths from 76 to 202 residues and includes monomeric, homodimeric systems and a ligand-bound complex. The restraint data included both unambiguous and ambiguous NOE-derived, dihedral and RDC restraints, ambiguous restraints, and a ligand-bound complex. This combination reflects realistic NMR structure-determination projects while remaining sufficiently concise to permit meaningful, interpretable comparisons of NEF read/write behaviour across independent software implementations.

Quaternary structure across the test set was deliberately varied. The test set included all-α, all-β and mixed α/β folds. While most entries are monomeric proteins, several homodimeric systems were included (2JR2, 2JUW, 2KKO, 2KO1) to assess correct handling of multi-chain assemblies. These homodimeric protein data sets also included X-filtered NOESY data for distinguishing interchain NOEs. In these cases, the structures comprise a single polymer entity represented by multiple polymer instances, providing a focused test of chain identifiers, atom referencing, and restraint assignment across symmetry-related subunits.

Additional restraint complexity was incorporated in a targeted manner. Residual dipolar coupling (RDC) data are present in three entries: 2KW5, which contains two independent RDC datasets, as well as 2KZN and 2LOY, enabling evaluation of NEF’s support for orientation-dependent restraints and associated metadata. Ambiguity handling was explicitly tested using a dataset containing ambiguous distance restraints (2PNG), while the remaining entries contain only unambiguous restraints. This stratified inclusion allowed ambiguity support to be assessed without making it a pervasive feature of the entire test set.

Using the NEF datasets, developers implemented and assessed their support for NEF through a round-robin exercise to test interoperability. Each developer assessed NEF read compliance, importing every dataset and running their standard workflows. Write interoperability was tested by exporting NEF from their software and having at least one other tool read and process it, principally using CcpNMR AnalysisStructure and the wwPDB OneDep validation suite (*20, 21*). We then worked with developers to identify and address any issues and re-tested until all issues were resolved and the specifications were finalized and standardized as Version 1.2 (<u>NEF/specification/v1_2_under_review at master · NMRExchangeFormat/NEF · GitHub</u>).

Several packages also exported specific restraints via private namespaces, e.g., MELD and UNRES introduced terms in their own namespaces, consistent with NEF’s extension mechanism, whereas ARIA includes additional project metadata as well as an additional saveframe with restraint violation data (Supplementary Fig. S3).

### NEF usage in restraint analysis

NMR-based structure determination involves multiple algorithmic steps, including iterative peak interpretation, ambiguity handling, restraint generation and refinement. Different software packages implement distinct strategies for interpreting identical experimental inputs, which may lead to differences in the resulting restraint sets and consequently, the structural ensembles obtained. The goal of the following analysis is not to benchmark or rank these algorithms, nor to assess absolute structure quality. Rather it is representative of the first round in the structure calculation process and demonstrates both in use and by analysis that NEF enables quantitative, restraint-level comparison of the interpretation of identical experimental data across independent workflows. By preserving explicit links between NOESY peaks, as in the current use case, and the restraints derived from them, NEF enables a researcher to assess how the input data are used during structure calculation, and to distinguish differences arising from algorithmic interpretation of the data versus those arising from differences in input formats.

Using the _`nef_restraint_links` saveframe enables the direct comparison of restraint sets generated by different programs from identical experimental data sets. We examined the restraints produced under two commonly used calculation conditions, i.e. using all programmatically detected peaks (denoted as unfiltered) and subsequently using a dataset in which the peaks were filtered on the basis of signal-to-noise (denoted as filtered), to assess how consistently peaks and restraints are interpreted by the three different structure calculation programs (Fig. 2). The workflow was tested for three experimental data sets, that were chosen as illustrative examples only, as the precise structural outcome is not the goal of this investigation.

**Figure 2.**
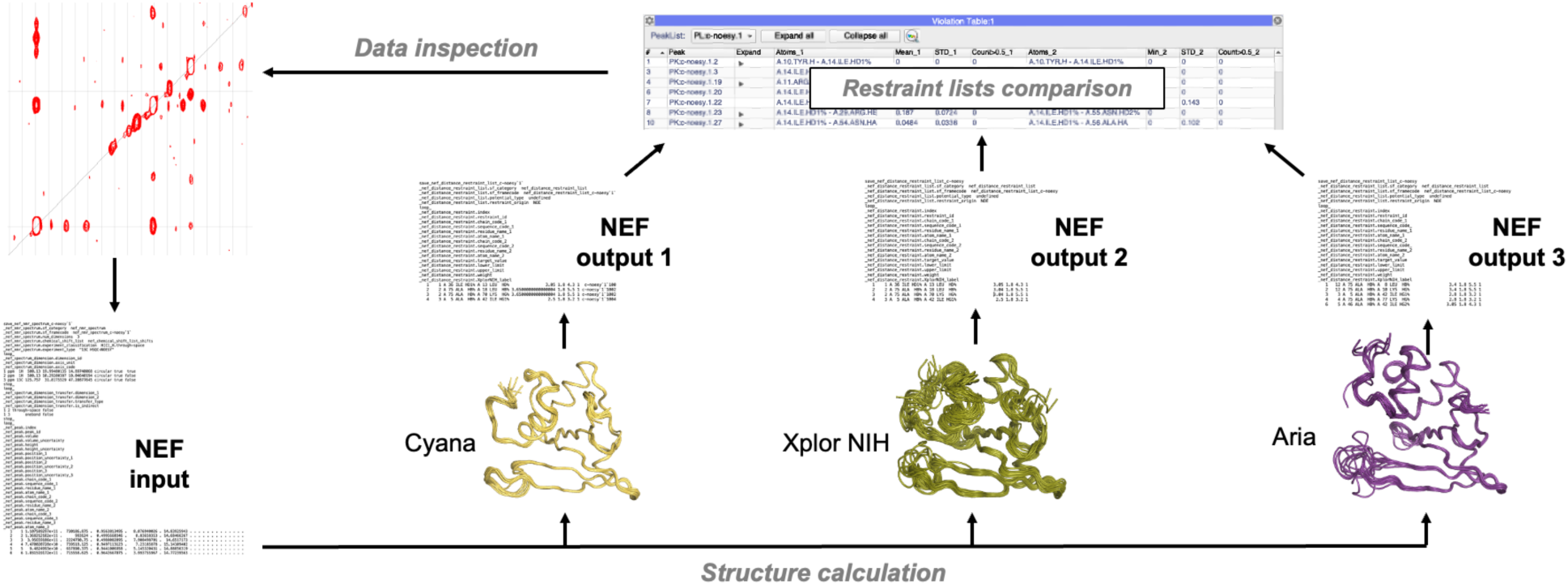
Workflow demonstrating NEF’s applicability across an extended NMR software ecosystem. NMR data (sequence, chemical shifts, peak lists and dihedral restraints) were exported as a NEF file, and processed by different structure calculation programs (ARIA, CYANA, and Xplor-NIH). Resulting structure ensembles and NEF files with restraint lists were compared to assess interoperability and consistency across the software packages.

#### Case 1: 2K3A –data rich protein

The 2K3A (*22*) dataset corresponds to the solution NMR structure of the *Staphylococcus saprophyticus* CHAP (cysteine-, histidine-dependent amidohydrolase/peptidase) domain, a 163-residue protein studied by the Northeast Structural Genomics Consortium (*23*). The dataset is characterised by high chemical-shift assignment completeness (∼93%), as reported in prior large-scale benchmarking studies (*24*), and by well-resolved and high-quality 3D ^15^N-NOESY and ^13^C-NOESY spectra. The high chemical-shift completeness reflects the quality of the upstream assignment work and provides a representative example of what can be achieved with high-quality protein datasets for which broadly similar behaviour across structure-calculation programs would normally be expected.

Using the unfiltered data with 5359 NOESY peaks, ARIA yielded restraints originating from 2746 peaks, compared to 2110 peaks used by Xplor-NIH and 1714 peaks used by CYANA. Here, the latter peak counts refer to the number of distinct input peaks that were converted into at least one distance restraint. All other peaks were not directly linked to a restraint, although they might have been interpreted, assigned and used by the program, e.g. for support in case of symmetric NOESY-peaks, or they might have yielded non-conformational restricting restraints which were not carried forward.

The associated ambiguity of the resulting restraints differs substantially between programs (24.2% for ARIA, 9.6% for Xplor-NIH and 7.2% for CYANA), reflecting ARIA’s greater tolerance of ambiguous assignments compared to CYANA’s more conservative treatment of peaks resulting in non-unique restraints.

Analysis by restraint type shows that all three programs unambiguously assigned a substantial number of long-range restraints (Figs 3A,4) when using the non-filtered peak lists, i.e. 626 restraints for ARIA, 421 restraints for Xplor-NIH and 524 restraints for CYANA, providing a strong basis for structure determination. Filtering the peak lists (S/N > 5) reduced the number of long-range atom-pair assignments derived from the filtered input in all three programs (Figs 3D,5), i.e. by 38.2% for ARIA, 24.4% for CYANA and 15.0% for Xplor-NIH, likely because long-range restraints more often result from weaker peaks. After filtering, the different programs agreed more closely on the peak to restraint assignments, as reflected by increased overlap between the restraint sets selected by each method (Figs 4,5). This is most evident for long-range restraints, for which the three-way unambiguous long-range overlap score, *i.e.* the number of identical long-range atom-pair assignments derived from the same input peaks in all three programs, is largely preserved under filtering, decreasing only slightly from 225 to 206 restraints (−8.4%).

**Figure 3.**
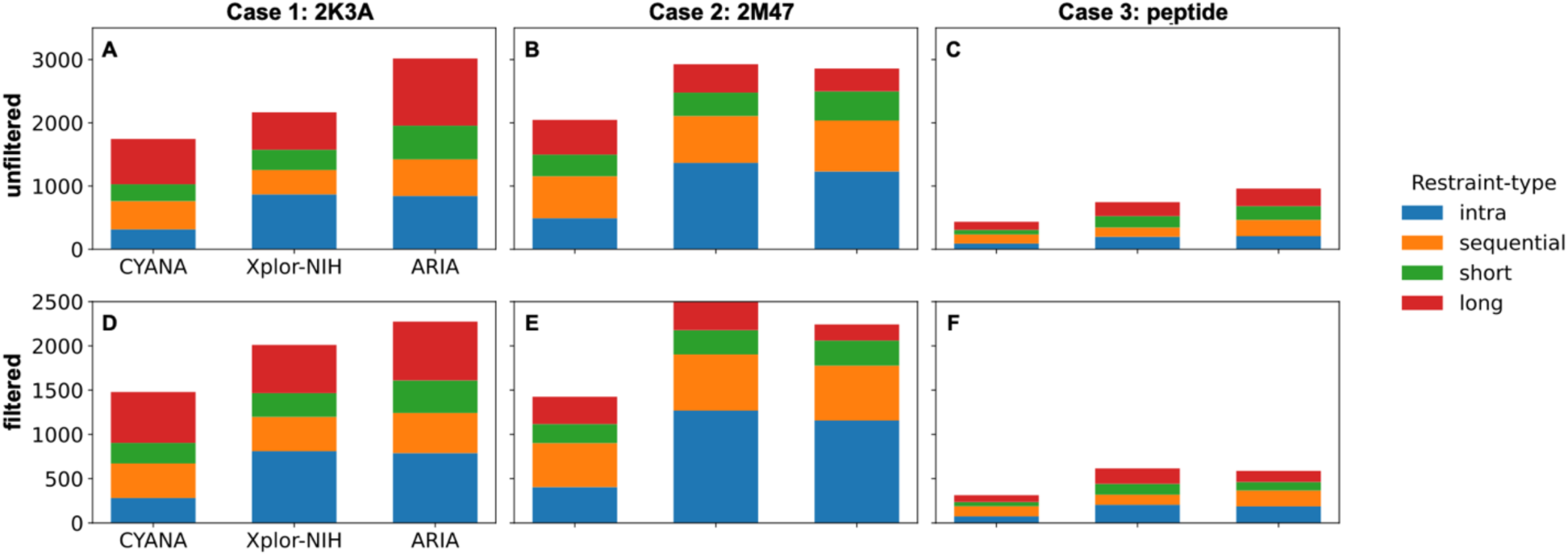
Restraint-type distributions across programs and noise conditions. Stacked bar charts show the number of restraints generated by ARIA, CYANA, and Xplor-NIH, stratified by NOE restraint type (intra-residue, sequential, medium-range spanning 2-4 residues, long-range). Columns correspond to individual programs, and panels are arranged by dataset (Case 1: 2K3A; Case 2: 2M47; Case 3: peptide). A-C) Results obtained using unfiltered peak dataset (i.e. all peaks included). D-F) Results obtained using a filtered peak dataset. Absolute counts are shown to emphasise retained experimental evidence rather than relative proportions and no correction for possible non-conformational restricting restraints has been made.

**Figure 4.**
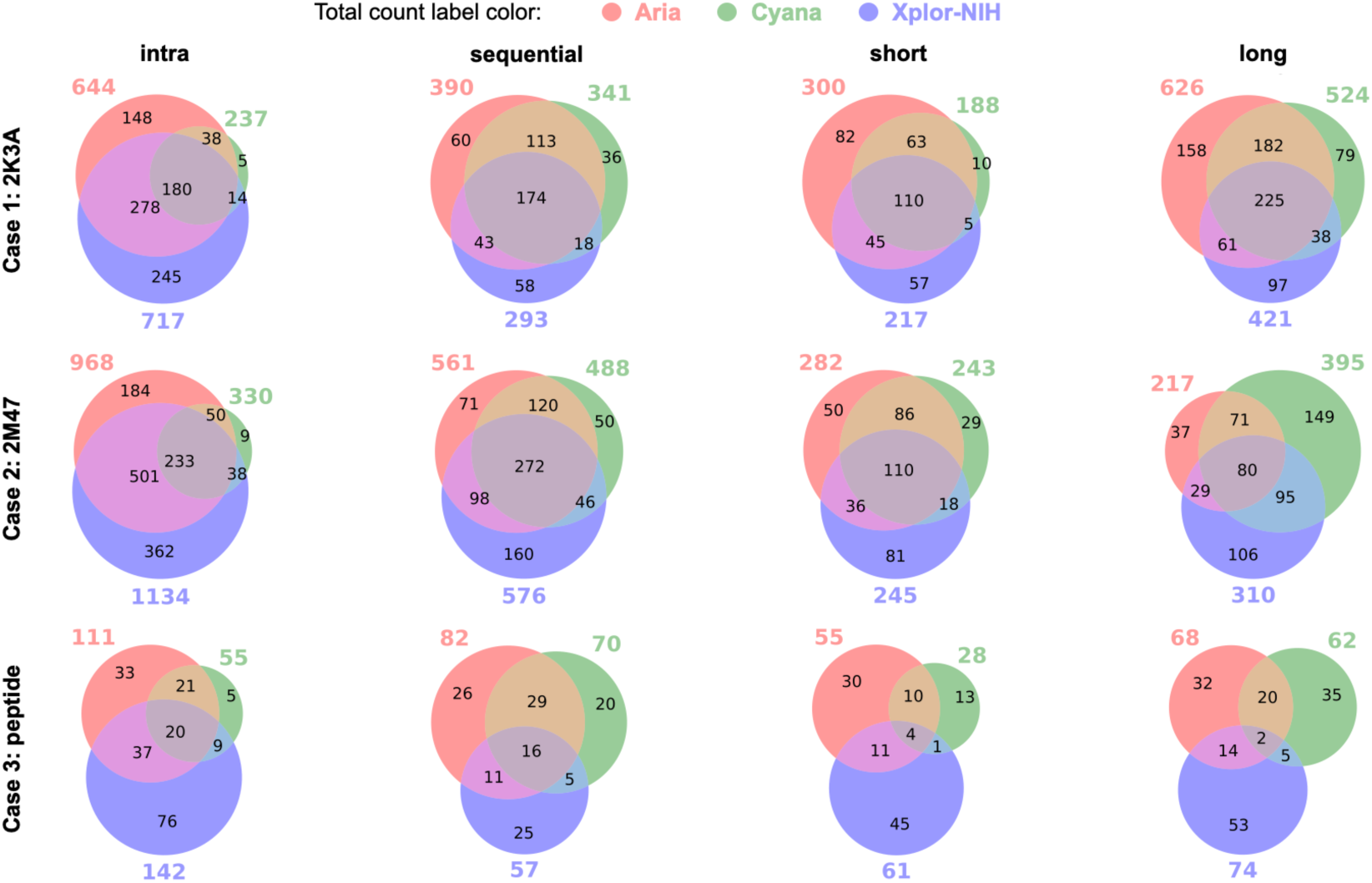
Restraint overlap across programs for unfiltered calculations. Multipaneled Venn diagrams summarising the overlap of distance restraints generated by ARIA, CYANA, and Xplor-NIH for calculations using unfiltered peak datasets. Rows correspond to datasets (2K3A, 2M47, peptide), and columns correspond to restraint-types (intra-residue, sequential, medium-range, long-range). Values indicate absolute numbers of unambiguous restraints. These diagrams highlight the size of the shared three-way core and method-specific restraint selection under identical experimental inputs.

**Figure 5.**
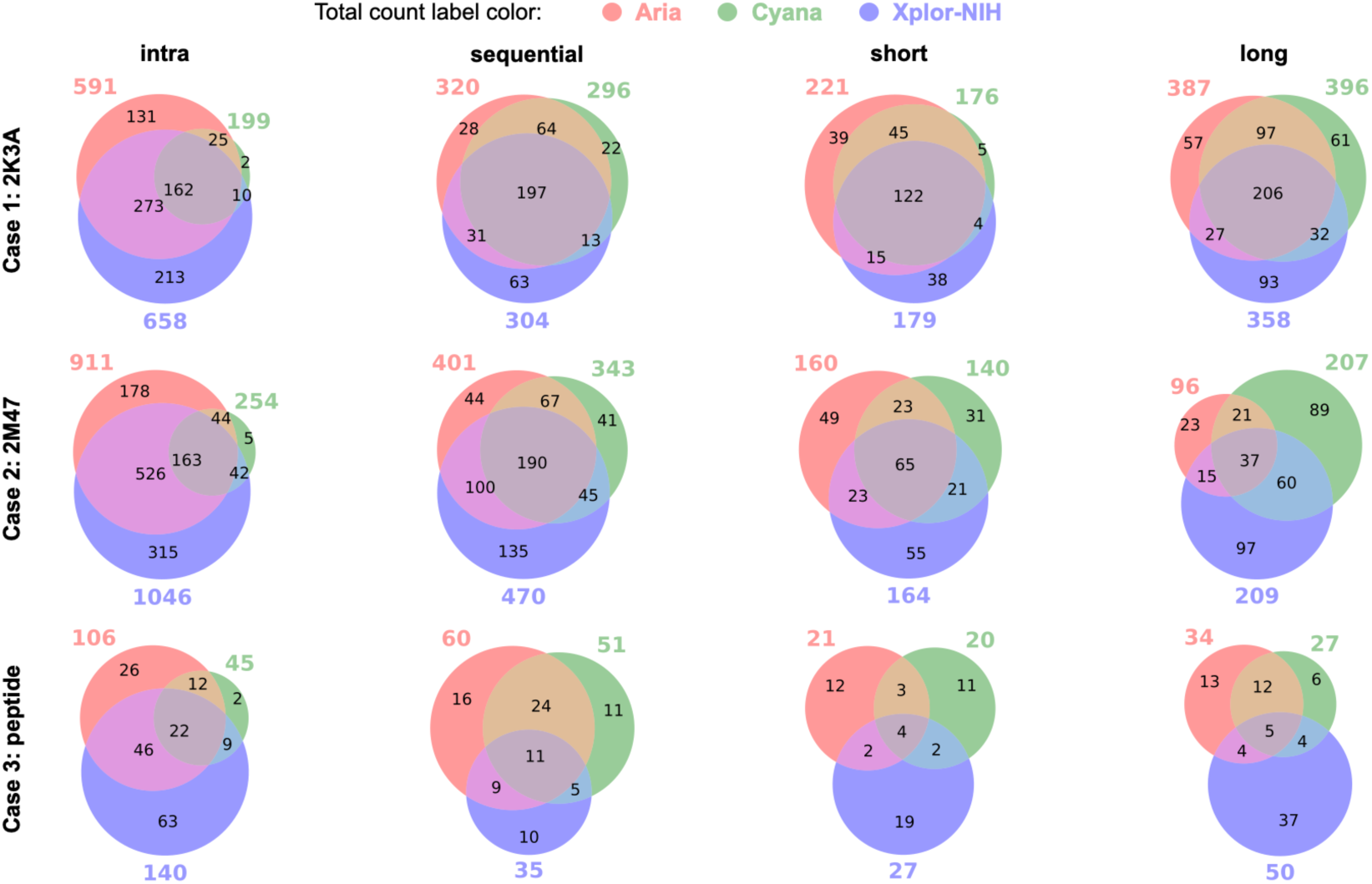
Restraint overlap across programs for filtered calculations. Venn diagrams equivalent to Fig. 4 but showing results obtained when using filtered peak datasets. Comparison with the results obtained using unfiltered data reveals how noise filtering alters both absolute restraint counts and the size of the shared three-way core, particularly for long-range restraints in noise-sensitive (Case 2) and underdetermined (Case 3) data sets.

The restraint-level observations are reflected directly in the calculated structure ensembles (Supplementary Fig. S4). For calculations using either non-filtered or filtered peak lists as input, CYANA, Xplor-NIH and ARIA all converge on the same global fold, with close agreement on secondary-structure elements and overall domain architecture. Differences between programs are limited to flexible loop regions and the termini, where modest ensemble dispersion is observed which does not affect the overall topology. Application of filtering does not induce any qualitative change in fold, consistent with the preservation of a shared core of long-range restraints supporting the same overall topology. Quantitative comparison to deposited structures or predictive models was not performed, as the purpose of this analysis is to examine consistency of experimental restraint interpretation rather than structural accuracy. Together, these results show that, for a well-behaved dataset, NEF enables consistent identification of a robust structural result across multiple programs, while also making systematic differences in restraint handling explicit.

#### Case 2: 2M47 – limited data protein

The 2M47 dataset corresponds to the solution NMR structure of the *Corynebacterium glutamicum* polyketide cyclase-like protein Cgl2372 (*25*), a 163-residue protein with 83% backbone and 89% overall chemical-shift completeness. Compared to the 2K3A case, this dataset exhibits increased spectral crowding and poorer signal to noise data, providing a realistic test of how different structure-calculation engines handle marginal or partially ambiguous experimental evidence.

Using the unfiltered data with 5891 NOESY peaks, Xplor-NIH generated restraints from 2835 peaks, compared with 2687 peaks for ARIA and 2010 peaks for CYANA. The corresponding fraction of ambiguous restraints, i.e. 10.3% for Xplor-NIH, 14.8% for ARIA and 13.2% for CYANA, indicate method-dependent differences in how ambiguity is tolerated and propagated when the data are less complete (Fig. 3).

Under filtering, the number of unambiguous long-range restraints was reduced substantially for all three programs (Fig. 5), with the strongest effect observed for ARIA (−55.8%), followed by CYANA (−47.6%) and Xplor-NIH (−32.6%). This behaviour highlights clear differences in how the programs trade off retaining a larger number of restraints versus prioritising only the most reliable long-range information under noisy conditions.

Overlap between unambiguous restraint sets increased modestly after filtering but remained consistently lower than in Case 1, reflecting greater divergence in restraint selection. Notably, the three-way unambiguous long-range score of 80 restraints when using the unfiltered dataset was already low compared to Case 1 but further reduces to only 37 restraints after filtering (−53.8%).

These changes at the restraint level have clear structural consequences (Supplementary Fig. S5). When all peaks are used, all three programs recover a broadly similar fold, but differences in ensemble dispersion are evident, particularly in the flexible loop regions. The ARIA ensemble encompasses a noticeably larger conformational space, whereas CYANA and Xplor-NIH generate more tightly clustered ensembles. After Filtering, these differences become even more pronounced: CYANA and Xplor-NIH maintain a recognisable core topology, while ARIA samples a larger conformational range. ARIA appears particularly sensitive to the removal of weaker, low-S/N peaks, consistent with the pronounced reduction in shared long-range restraints identified by analysis using the peaks, restraints and their links documented in the NEF files.

#### Case 3: short peptide – underdetermined regime

We used the NOESY data recorded for a 27-residue peptide of the N-terminal region of Neisseria gonorrhoeae TfpC (*26*), to establish how data provenance using NEF could aid the initial stages of a structural analysis of an underdetermined system with likely intrinsic conformational variability. The unfiltered data contained 1144 NOESY peaks derived from a single 2D NOESY spectrum with relatively challenging data quality. Using the unfiltered data, ARIA used 708 peaks (50.0% ambiguous), compared with 416 peaks for CYANA (9.6% ambiguous) and 621 peaks for Xplor-NIH (29.6% ambiguous). These differences reflect how each program handles ambiguity and uses peaks when experimental information is extremely limited.

Despite a substantial number of restraints defined by all three programs, the overlap between the derived restraints is limited. Only 39 restraints form a strict three-way consensus, defined as restraints derived from the same input peak with identical atom-pair assignments in all three programs, which increases to 93 if this definition is relaxed to entail at least one atom-pair assignment in common across all programs. By contrast, 846 peaks are assigned by at least one program but show no common atom-pair assignment across all three methods (Fig. 4). Using the filtered data reduces the total number of peaks and their conversion into restraints but does not qualitatively change this picture: overlap between programs remains low, and the shared core remains small relative to the total number of derived restraints (Fig. 5).

Unsurprisingly given the limited restraint overlap, the CYANA, Xplor-NIH and ARIA calculations do not converge on a common structure (Supplementary Fig. S6), highlighting how in underdetermined systems different algorithms may extract distinct, but internally consistent solutions from the same sparse and challenging experimental data. For reference, the AlphaFold3 () model suggests a compact β-hairpin conformation, including the possibility of a di-sulfide bond, which was not included in the current set of calculations.

### Adoption of NEF

NEF has become a standard for consistent, interoperable (FAIR) NMR data exchange across diverse software platforms. Its integration spans spectral analysis, assignment, structure calculation, validation and deposition (Fig. 6), and streamlines the transfer of chemical shift, peak and restraint data across these various stages. Table 1 lists current support among major programs and data repositories. Below, we will briefly discuss how individual packages implement NEF and the specific roles it plays within their workflows.

**Figure 6.**
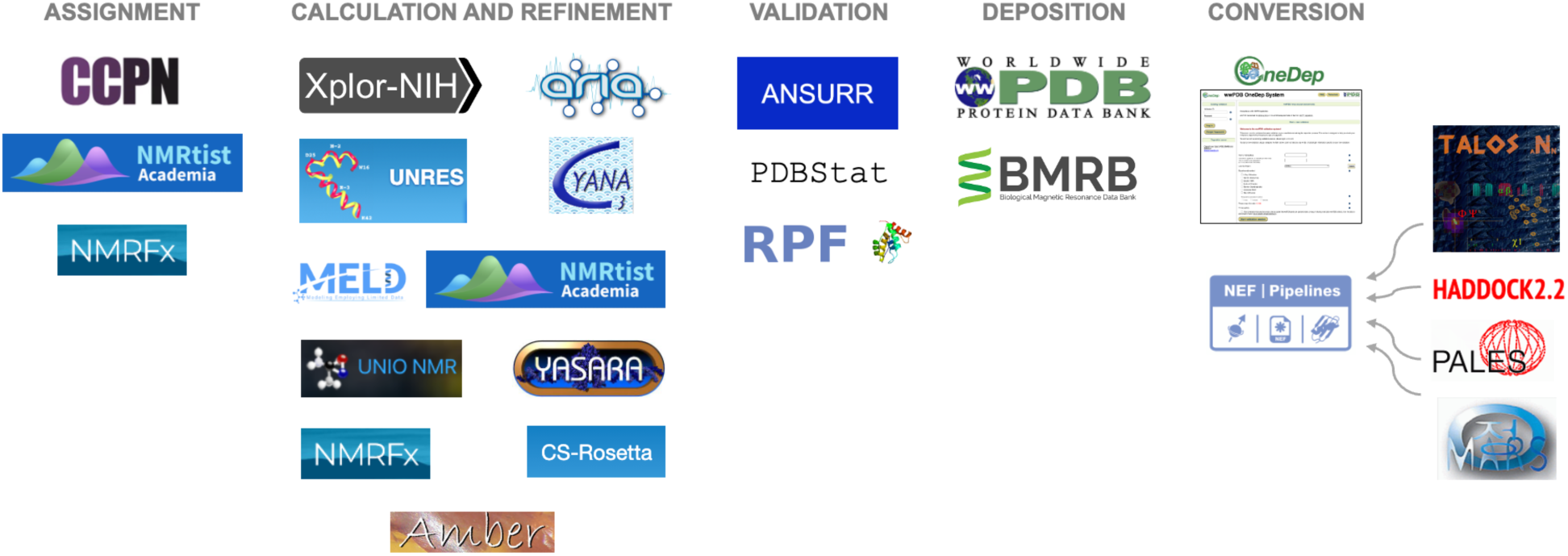
Adoption of NEF for integrated NMR data analysis. NEF enables the seamless transfer of crucial NMR data across pipelines for resonance assignment, structure calculation, refinement, validation, and deposition. Compatible software packages at each stage are using NEF-formatted data to drive analysis. Structures may also be sourced from predictions, for example from AlphaFold, with subsequent refinement using NEF-compatible pipelines.

Since 2024 NEF is used by both the BMRB (*27*) and the wwPDB (*28*) within its OneDep (*21*) system to support joint deposition of NMR data and derived restraints, alongside structural coordinates in mmCIF format. This coordinated NEF/mmCIF framework provides a common and structured format that enhances interoperability and facilitates the consistent deposition, validation and retrieval of NMR-derived structural information in public databases. By linking experimental provenance directly to structural models, NEF enables these organisations to manage and share complex NMR datasets more transparently, promoting consistency and accessibility within the structural biology community. Integration of NEF with deposition and validation (*29*) systems such as BMRB and wwPDB therefore supports a streamlined and unified workflow for data submission and archiving (*30*).

The CcpNmr Analysis software was an early adopter of the NEF standard (*31, 32*). It was used in this work for setting up structure calculations using Xplor-NIH, CYANA and ARIA (*vide infra*) and the subsequent import of their NEF output into CcpNmr AnalysisStructure. NEF serves as the common data standard for different structure calculation programs, facilitating workflows that provide users with the flexibility to choose the most suitable structure calculation program while maintaining compatibility and ease of comparison. A variety of tools have been implemented to analyse and compare restraints from different structure calculation runs as used for the analyses shown above.

Xplor-NIH (*18, 33*) is a widely used software package for macromolecular structure determination that combines experimental NMR restraints with molecular mechanics and simulated annealing protocols. Support for NEF within Xplor-NIH enables the direct import and export of restraint data in a standardised format, reducing ambiguity associated with atom naming and restraint representation.

ARIAweb (*15*) is a web server designed to improve the accessibility and usability of the ARIA (Ambiguous Restraints for Iterative Assignment) tool (*34*) using CNS (*35*) for structure generation, and providing an online platform for NMR structure calculation projects (*15*). ARIAweb has adopted the NEF standard to integrate diverse data types to simplify automated NOE assignments.

CYANA (*17*) is an established program for automated chemical shift assignment and NMR structure calculation that combines iterative NOE assignment with rapid structure generation using torsion-angle dynamics. By including CYANA command scripts in the metadata saveframe, NEF files can be made executable by CYANA, which combines the data and the commands for an assignment or structure calculation in a single file.

ARTINA (*36*), available at the NMRtist webserver (*37*), is a machine learning approach that covers the complete workflow from spectrum analysis to assignment and structure calculation without human intervention. NEF is supported for the input and output of chemical shifts, peak lists and restraints.

Beyond these core structure determination engines, NEF also enables integration with tools that emphasise automation, modelling, or simulation. NMRFx (*38*) is another application that was an early adopter of NEF. It incorporates NEF within an end-to-end processing, assignment, shift prediction and structure calculation environment, allowing for complete analysis and structure calculation within the software, or the use of NEF to interact with other NEF supporting applications. UNIO-NMR (*39*) uses NEF-compatible representations within automated pipelines to support reproducible calculations with minimal manual intervention. CS-Rosetta (*40*) uses NEF- encoded chemical shifts and restraints to guide fragment-based modelling, allowing comparison between NMR-driven and modelling-centric structure generation strategies.

Finally, NEF provides a route for incorporating experimental NMR information into physics-based simulation frameworks such as UNRES (*41*), MELD (*42*), YASARA (*43*), and AMBER (*44, 45*), where restraints are mapped onto molecular dynamics–based sampling and refinement protocols. In this broader context, tools such as NEF-Pipelines (https://github.com/varioustoxins/NEF-Pipelines) facilitate the transfer and manipulation of NMR data and NEF files themselves, including peaks, chemical shifts, and sequence information, while in transport between a wide range of NMR programs, including legacy software that may never become natively NEF-compatible. Similarly, PDBStat supports NEF-centred workflows by enabling reliable interconversion between NEF and legacy formats and by providing restraint analysis and structure quality assessment to support validation and consensus analysis across methods (*46*). PDBStat also provides an interface between NEF and programs not yet specifically modified to use NEF-formatted restraints such as AutoStructure / ASDP (*47, 48*) and NMR-restrained Rosetta.

While these approaches focus primarily on biomolecular structure determination and validation, the same NEF-based principles can be applied to other NMR applications that rely on consistent, time-resolved experimental data representation.

#### Ligand Screening

CcpNmr AnalysisScreen^26^ is a platform for ligand screening workflows which enables efficient analysis of small-molecule NMR data across a wide range of experiments, including ¹H, ¹⁹F, STD, Water-LOGSY and CPMG. By storing screening data in NEF, time-dependent, chemistry-induced changes across ligand libraries can be tracked explicitly and reproducibly, yielding improved sensitivity and reliability in the identification of potential binders (Mureddu *et al.*, manuscript in preparation).

#### Metabolomics

NMR-based metabolomics is a rapidly growing field, valued for its reproducible, non-destructive measurements and minimal sample preparation. To support metabolomics research, the CcpNmr Analysis Simulated Metabolomics DataBase (CASMDB) (*49*) aggregates curated metabolite information from the Human Metabolome Database (HMDB) (*50, 51*), the BMRB, and GISSMO (*52*). CASMDB provides simulated spectra with associated metadata for a broad range of small molecules, enabling compound identification, method development, and benchmarking within a consistent and reproducible framework. All data are stored in NEF, ensuring explicit links between chemical shifts, spectra, and compound metadata, and enabling interoperability with NEF-compliant software such as CcpNmr AnalysisMetabolomics (*53*).

## Discussion

We defined NEF as a general standard for NMR data exchange, providing unambiguous data definitions for chemical shifts, peaks and various restraint types, accompanied by crucial metadata to allow for FAIR data handling (Fig. 1, Supplementary Figs S1, S3). Community-led interoperability testing established the integrity of the data after reading of NEF files, and where applicable, also the writing of NEF files. This establishes NEF as a reliable foundation for inter-software data exchange, enabling data analysis workflows across multiple software packages (Fig. 6).

The development of NEF was initially driven by the need for standardized and unified NMR data deposition in the wwPDB (*28*), linking structural data to their underpinning experimental data, thus allowing for proper NMR structure validation pipelines (*29*) at the point of deposition. However, NEF can be used not only to move data between programs, but also to enable clear, restraint-level comparison across different assignment and structure-calculation workflows using the same experimental data or to allow for comparison of a variety of input data. NEF enables these comparisons to be performed without the need to adapt existing computational protocols or enforce artificial uniformity on downstream algorithms. Each program operates according to its native logic, yet the resulting restraints can be interrogated within a common framework because their provenance and linkage to experimental peaks are preserved.

We tested the NEF using three automated NOE assignment and structure calculation engines and three datasets (Fig. 2), that span a realistic range of experimental regimes, from a well-behaved protein, through a noise-sensitive case, to an intrinsically underdetermined and challenging peptide, which reflect the range of possibilities encountered in NMR-based structure determination. These analyses illustrate how NEF enables transparent comparison of the experimental evidence used by each workflow. Overlap metrics do not in themselves establish structural correctness; rather, they quantify the degree to which independent algorithms can identify a shared core of experimental support from identical input data such as chemical shifts and peak lists. In well-determined systems, this shared core remains robust under data filtering, whereas in underdetermined regimes it collapses, revealing the extent to which structural outcomes depend on algorithmic interpretation rather than convergent experimental evidence.

The analyses nevertheless reveal several qualitative behaviours that are consistent with prior observations in NMR structure determination, even though their detailed investigation lies beyond the scope of the present work. First, inclusion of weaker peaks frequently increases the number of retained long-range restraints and can improve apparent structural convergence, highlighting the importance of redundancy in experimental evidence (*54*). The impact of the quality of the peak data was also identified previously by the CASD-NMR-2013 study (*5*), albeit at a per-residue level whereas in this study we can compare the interpretation of the individual peaks. Second, ensembles of similar structural precision can arise from substantially different restraint sets or interpretation regimes, indicating that apparent precision does not necessarily reflect a uniquely supported experimental solution. Finally, differences in structural between programs largely reflect distinct tolerances to ambiguity and sparse data, rather than differences in underlying experimental input. These observations emphasise that interpretation of NMR data depends not only on the available measurements, but also on how algorithms manage uncertainty and redundancy. A systematic investigation of these effects, including sensitivity to stochastic sampling and minimal versus redundant restraint representations, has been explored previously (*55, 56*) and will be addressed in future work.

Across the three cases, the comparison reveals clear patterns (Figs 3-5), although they do depend on the dataset. In Case 1 (2K3A, unfiltered peaks), ARIA retains the largest restraint set, driven by higher ambiguity retention, while CYANA is the most conservative. In Case 2 (2M47, unfiltered peaks), Xplor-NIH retains the largest set, with ARIA intermediate and CYANA again the most conservative. Across all cases and conditions, ARIA retains a higher fraction of ambiguous restraints, CYANA applies the strongest filtering, and filtering increases pairwise agreement.

The three-way unambiguous long-range overlap score (Figs 4,5) provides an illustrative measure of data consistency that becomes accessible when experimental provenance is preserved via NEF. Importantly, this analysis is not intended to imply that multiple structure-calculation programs must be applied routinely. Instead, it demonstrates how NEF enables independent workflows to be compared at the level of experimental evidence when such comparisons are desired. For well-behaved datasets, the shared restraint core remains largely stable under peak filtering, whereas in more challenging or underdetermined regimes it collapses. Similar behaviour has previously been linked to stochastic sampling effects and variability between calculation runs, where apparently precise ensembles may arise despite limited consensus experimental support (*57*). Our analysis of Case 3 illustrates how such underdetermination can become visible when restraint provenance is explicitly traceable.

The comparison of unfiltered and filtered data illustrates another practical advantage: transformations to the data applied upstream or downstream of the structure calculation remain traceable. Noise filtering consistently reduces absolute restraint counts while increasing relative agreement between methods, a trade-off that would be difficult to characterise without explicit peak-to-restraint linkage.

From a workflow perspective, the results demonstrate that NEF supports cross-software, multi-iteration analysis (Fig. 6) within a single project, rather than enforcing reliance on a single structure-calculation engine. Restraints generated by different programs or by the same program under different conditions can easily be compared and rationalised using a shared representation of the available experimental data, enabling genuine loss of experimental evidence to be distinguished from algorithmic reinterpretation of identical data. This distinction is particularly important when diagnosing divergent structure calculations or when selecting tools for specific stages of an iterative workflow. NEF-based protocols therefore shift evaluation from qualitative descriptions of “agreement” or “convergence” to quantitative assessment of shared experimental support.

Note that although we have illustrated these advantages of NEF over software-specific formats in the context of NMR structure calculation, they apply equally to tracing the origin of chemical shift assignments, or comparing chemical shift assignments determined by different manual/semi-automated (*58*) or automated (*36, 59*) methods, and even in integrative deep learning approaches using AlphaFold and chemical shift prediction (*60*).

### NEF future directions

Looking ahead, the future of NEF will see existing gaps addressed, improving its flexibility for users and software developers. Below we discuss two pertinent topics.

#### Topology

The current NEF implementation captures basic topological information through the `nef_molecular_system` saveframe, which lists chains, residues, atoms and bonds in a standardized form. Within this framework a “residue” may represent any molecular component, including ligands and ions. Only essential information describing covalent bonds beyond the standard NEF residue definitions is recorded in the molecular system saveframe. This enables consistent interpretation of restraint and chemical shift data across software platforms. To maintain NEF’s role as an NMR-data exchange format, its definition deliberately excludes detailed molecular mechanics parameters, such as bond lengths, charges, bond angles and improper terms, that are primarily required for molecular dynamics simulations. Future assessments will also include validation of models against NEF-formatted NOESY peak lists (*61*).

Future iterations of NEF could also integrate links to external chemical component dictionaries, e.g. as provided by CcpNmr or wwPDB’s Chemical Compound Dictionary (*62*), to ensure accurate atom typing and facilitate cross-referencing with other structural biology data resources. Another area for development is the representation of ensemble or time-dependent topologies, allowing NEF to describe systems where connectivity changes over time, such as during chemical reactions, ligand binding, or conformational rearrangements.

#### Dynamics

The current NEF specification provides robust support for chemical shift assignments, peaks, and restraints. A next stage of NEF development is its much-needed extension to include NMR dynamics data. An expansion of NEF that carries NMR dynamics data and associated metadata from spectra to derived parameters, while preserving provenance has been proposed (*63*). Concretely, NEF should pair a compact table of observables, e.g. the measured series of intensities or ratios across one or multiple spectra indexed by the experimental axis, such as time, CPMG repetition rate, offset, or B₁, with a companion table of derived results, such as R1, R_2_, ¹H–¹⁵N NOE, R_ex_, k_ex_, Δω, model name, parameter values with uncertainties, and goodness-of-fit. Stable identifiers link peaks to series points to derived results on a per residue or per-atom basis, akin to the restraint links saveframe, so that refitting, auditing and downstream reuse become routine rather than the exception.

#### Modelling with other kinds of NMR data

The NMR Exchange Format (NEF) establishes a unified and extensible framework for representing biomolecular NMR data that bridges the persistent gap between software-specific data models. Through direct testing, we demonstrate that NEF enables consistent, lossless exchange of restraints, chemical shifts and spectral metadata, across multiple software packages, a key milestone toward reproducible NMR data analysis, structure determination and structure validation. Its adoption by major software tools and data repositories, including CcpNmr, ARIA, CYANA, Xplor-NIH, BMRB and the wwPDB, now supports a strategy where tools work together for NMR data analysis and deposition. By directly linking experimental observations to derived restraints, NEF ensures data traceability from peak list to final structural model.

Future development of the NEF schema should also accommodate paramagnetic relaxation enhancements (PREs), pseudo-contact shifts (PCSs), chemical shift perturbations (CSPs) or any related series where the experimental axis may be distance, orientation, frequency or concentration. These datasets all benefit from the same pattern: capture the observable series with units and axis details; link to a derived result with named parameters, e.g., χ tensor terms for PCS, binding constants for titrations, etc. and record the uncertainty model and goodness-of-fit. Where experiments require extra context NEF provides optional, clearly named fields so that interpretation is unambiguous without introducing complexity.

The implications extend well beyond structure determination. NEF’s structured, FAIR-centred design allows for better integration of NMR data into wider data analysis pipelines such as those implemented for ligand screening and metabolomics, as well as artificial intelligence and machine learning approaches. Its open, extensible design provides a way to accommodate new data types, from dynamic measurements to hybrid structural techniques, without compromising compatibility.

Its ability to cross-reference other database resources allows for usage of NEF in integrated structural biology approaches because references to relevant other data, e.g. gene information, EM maps, SAXS/SANS data, can be carried along.

In summary, NEF builds on and extends earlier efforts such as the NMR-STAR (*1*) or CcpNmr data model (*2*), by providing a lightweight, yet comprehensive framework specifically designed for the interoperable exchange of NMR data. Adoption by fifteen programs or software packages transforms the NMR data from information contained in isolated, program-specific files into interoperable scientific resources that can be used across a wide array of analyses. We hope that as adoption continues, NEF will catalyse more transparent, collaborative, and reproducible workflows across structural biology, strengthening NMR’s role as a technique for integrative biomolecular research.

## Methods

### NEF

NEF development is hosted on its GitHub site (https://github.com/NMRExchangeFormat/NEF). A fully annotated example NEF file is available at the repository. Data and scripts used for this manuscript are also available on GitHub as part of the NEF development framework. The NEF charter governs its functioning as a community-driven effort, including future changes and extensions.

### NEF testing Datasets

Fifteen well-curated datasets (Supplementary Table ST1) were produced using standardized experimental and analysis pipelines. Each set contained NEF file, which included all mandatory saveframe (metadata, sequence and chemical shift list) and restraints (distance, and where appropriate dihedrals and RDC). Structure coordinates in mmCIF format was available along NEF file. These provide a robust baseline for interoperability testing. They were chosen to cover a broad range including sparse data, a protein-ligand complex and homodimers. In addition to distance- and dihedral angle restraints, some datasets also included residual dipolar couplings (RDCs). Except for one protein-ligand complex, all test cases involve only canonical amino acids, with no post-translational modifications or non-standard residues. This ensured that differences observed during data exchange could be attributed to format handling, atom naming, restraint representation, and metadata consistency, rather than to topology or chemistry-specific issues outside the current scope of NEF. The protein-ligand complex (6NBN) allowed software packages to test handling of such systems.

The NEF files for the test datasets were all created from NMR-STAR files deposited in the BMRB. NEF adopts IUPAC-based nomenclature with extensions for prochiral atom assignments, whereas the BMRB’s NMR-STAR format uses ambiguity codes to describe atoms and atom groups in a way that aligns with these principles. We observed several instances of inconsistencies with the BMRB ambiguity codes, likely because this information is supplied by depositors and is not subject to detailed curation During the conversion from NMR-STAR to NEF, particular attention was paid to identifying and correcting such discrepancies while preserving the original input when appropriate. In cases where the ambiguity could not be resolved with confidence, atoms and atom groups were deliberately left stereo-specifically assigned or changed to non-stereospecific assignment depending on contextual information. This approach helped to maintain consistent annotation within the NEF files used for testing.

### Application of NEF

Three examples were chosen for testing the NEF-based structure calculation protocols using actual experimental data, representing data-rich (wwPDB entry 2K3A, NESG-id SyR11), data-limited (wwPDB entry 2M47, NSGC-id CgR160) and a short highly underdetermined 27 amino-acid peptide of the N-terminal region of *Neisseria gonorrhoeae* TfpC (*26*). For all three data sets, experimental NMR data and assignments were available (Supplementary Table ST2). For Case-1 and Case-2, prior structural results have also been published.

### Peak picking and preprocessing

Peak picking was performed using CcpNmr AnalysisStructure v3.2.2 (*31*) with default parameters for each spectrum type, using the data as supplied. To ensure consistency for two protein datasets the noise level was estimated using the program’s built-in noise-calculation tool in an area of the spectrum not expected to have any peaks and contours were dropped an additional 10% lower to provide extensive peak coverage. This intentionally permissive threshold was chosen to retain weak but potentially informative signals, ensuring that differences observed between programs arise from downstream interpretation rather than from upstream peak-selection choices. No manual inspection, curation, or post-processing of peak lists was carried out. Synthetic 2D ^1^H-^13^C-HSQC and ^1^H-^15^N-HSQC peak lists derived from the assigned chemical shifts were used to guide peak picking in the corresponding 3D ^13^C-NOESY and ^15^N-NOESY spectra, providing a consistent reference for peak identification across two protein datasets. For the peptide, 2D NOESY peaks were picked in regions corresponding to chemical shift coverage. In each case, peak lists were generated once and reused unchanged across all structure-calculation programs.

To assess robustness to noise, an additional peak list was generated for each spectrum by applying a simple signal-to-noise threshold (S/N > 5). These two peak sets are denoted as unfiltered or filtered, respectively, with the latter representing commonly used filtering choices in NMR-based structure determination. The unfiltered and filtered data sets were treated as independent inputs for downstream calculations.

### Ab initio structure calculations from experimental data

*Ab initio* structure calculations were performed using the default protocols implemented in CYANA (*57*), and Xplor-NIH (*33, 64*) and a slightly modified protocol for ARIA/CNS (*15*) (20,000 cooling steps, 30 calculated and 10 used structures per iteration, restraint combination for the first four iterations and selecting the log-harmonic potential (*65*)) with all programs using identical experimental inputs of either unfiltered or filtered data provided via NEF files. Distance restraints were derived automatically from NOESY peak lists using each program’s native assignment and calibration procedures and subsequently used within each program’s native assignment, calibration procedures and each software package’s own structure generation protocol, including its internal geometry definitions and optimisation strategy,

Each program interpreted the data independently generating final structure ensembles of between 10-20 conformers and their associated restraints, which were written as NEF files and preserving explicit links between experimental peaks and derived restraints. For CYANA a _`nef_peak_restraint_links` saveframe was added as part of post processing. These outputs formed the basis for the restraint-level analyses reported in the Results section.

### Restraint extraction and classification

Restraint lists in NEF format were imported into CcpNmr AnalysisStructure for comparison, enabling direct, restraint-level comparison across the originating programs without additional file conversion or loss of provenance. Restraints were grouped by originating program and classified according to restraint-type, i.e. intra-residue, sequential, medium-range (spanning 2-4 residues), or long-range as advocated by the wwPDB NMRvalidation taskforce recommendation (*66*) and ambiguity status (ambiguous or unambiguous), based on the number of alternative atom identifier pairs associated with each restraint identifier.

Explicit links between peaks and restraints, recorded in the _`nef_peak_restraint_links` saveframe, were used to ensure that all derived restraints could be traced back to their originating peaks. The mapping between peaks and restraints is not strictly one-to-one. A single NOESY peak may correspond to multiple candidate atom pairs when assignments are ambiguous, resulting in multiple alternative restraints derived from the same peak. Conversely, multiple peaks may support the same atom pair—for example through redundancy across spectra, symmetry-related signals, or program-specific merging and calibration procedures—so that several peaks may contribute to a single final restraint. Peaks may also contribute restraints spanning multiple classes; for example, a single peak can be ambiguously associated with both short-range and long-range contacts.

### Restraint overlap analysis and metrics

Restraint overlap between programs was assessed using Venn region counts based on exact matching of atom-pair assignments. For each case and peak list condition, restraints were classified as unique to a single program, shared between pairs of programs, or shared across all three. Absolute restraint counts were computed directly from these classifications.

Pairwise agreement between restraint sets was assessed using direct overlap counts derived from set intersections and unions, reported as percentages where appropriate.

### Visualisation and reporting

Results were visualised using stacked bar charts to show absolute restraint counts by program and restraint-type, and Venn diagrams to summarise shared and program-specific restraint sets. All figures were generated using Matplotlib (*67*) from intermediate tabulated data exported alongside the plots to ensure reproducibility and auditability.

## Acknowledgements

GWV, BOS, GT acknowledge support by UKRI/MRC (grant MR/V000950/1). GWV acknowledges support by STFC/CoSeC for the DRIIMB project.

CDS was supported by the Intramural Research Program of the National Institute of Diabetes and Digestive and Kidney Diseases, National Institutes of Health.

BMRB is currently supported by NIH grant R24GM150793. GTM was supported for this work by NIH grant 1R35GM141818.

RCSB PDB core operations are jointly funded by National Science Foundation (NSF) (DBI-2321666, PI: S.K. Burley), the US Department of Energy (DE-SC0019749, PI: S.K. Burley), and the National Cancer Institute, the National Institute of Allergy and Infectious Diseases, and the National Institute of General Medical Sciences of the NIH (R01GM157729, PI: S.K. Burley)

SR and JG were supported by UKRI/MRC (grant MR/W000814/1).

AL and EL acknowledge financial support from the National Science Centre, Poland (grant UMO-2023/51/B/ST4/01218), the use of supercomputing resources at the the Centre of Informatics—Tricity Academic Supercomputer and Network (CI TASK) in Gdańsk, at the Interdisciplinary Center of Mathematical and Computer Modeling (ICM), the University of Warsaw, under Grant No. GA71-23, and the local 796-processor Beowulf cluster at the Faculty of Chemistry, University of Gdańsk.

PG acknowledges support from the Japan Society for the Promotion of Science (JSPS) Grant-in-Aid for Scientific Research (23K05660).

## Author contributions

EP, GWV and GTM designed the research; GWV, RHF and AG developed the NEF standard; EP, RT and KB assembled the testing data; EP, KB, RT and all program authors performed the testing; EP, BB and CDS performed the structure calculations; JY led the wwPDB projects on incorporating NEF into OneDep deposition and validation system; Ezra P. developed PDBx/mmCIF data model for NEF-mmCIF conversion; EP and GWV wrote the manuscript; All authors read and approved the text.

