## Supplementary Materials for "The NMR Exchange Format (NEF): Specification and Applications"

Eliza Płoskoń[1,*], Kumaran Baskaran[2,*], Roberto Tejero[3,*], Charles D. Schwieters[4], Benjamin Bardiaux[5,6], Peter Guentert[7,8,9], Rasmus H. Fogh[1,10], Aleksandras Gutmanas[11,12], Edward J. Brooksbank[1], Masashi Yokochi[13], David S Wishart[14], Jonathan R Wedell[2], Wim F. Vranken[15,16] Daniel Thompson[1], Gary Thomson[17], Brian O. Smith[18], Saima Rehman[19], Theresa A. Ramelot[20], Timothy J. Ragan[1], Alberto Perez[21], Binod L. Perera[21], Ezra Peisach[22], Michael Nilges[6], Luca G. Mureddu[1], Arup Mondal[21], Emilia A. Lubecka[23], Adam Liwo[24], Genji Kurisu[13], Naohiro Kobayashi[13], Piotr Klukowski[7], Bruce A. Johnson[25], Yuanpeng J. Huang[20], Jeffrey C. Hoch[2], Victoria A. Higman[1], Torsten Herrmann[26], Morgan W. Hayward[1], James A. Garnett[19], David A. Case[27], Stephen K. Burley[22], Paul D Adams[28], Gaetano T. Montelione[20,‡], Geerten W. Vuister[1,‡]

1. Division of Molecular and Cellular Biology, Leicester Institute of Structural and Chemical Biology, University of Leicester, Lancaster Road, Leicester, LE1 9HN, United Kingdom.
2. Biological Magnetic Resonance Data Bank, Department of Molecular Biology and Biophysics, UCONN Health, Farmington, CT-06030, USA.
3. Departamento Química Física. Universidad de Valencia. Avda Dr. Moliner, 50. Burjassot (Valencia). Spain.
4. Laboratory of Chemical Physics, National Institute of Diabetes and Digestive and Kidney Diseases, National Institutes of Health, Bethesda, Maryland, USA.
5. Bacterial Transmembrane Systems Unit, Institut Pasteur, Université Paris-Cité, CNRS UMR3528, 75015, Paris, France.
6. Structural Bioinformatics Unit, Institut Pasteur, Université Paris-Cité, CNRS UMR3528, 75015, Paris, France.
7. Institute of Molecular Physical Science, ETH Zurich, Zurich, Switzerland.
8. Institute of Biophysical Chemistry, Goethe University Frankfurt, Frankfurt am Main, Germany
9. Department of Chemistry, Tokyo Metropolitan University, Hachioji, Tokyo, Japan
10. Current address: Global Phasing Ltd., The Quad 9, Journey Campus, Castle Park, CB3 0AX, Cambridge, UK.
11. Protein Data Bank in Europe, European Molecular Biology Laboratory, European Bioinformatics Institute (EMBL-EBI), Wellcome Genome Campus, Hinxton, Cambridge, CB10 1SD, UK
12. Current address: Data Science and AI, Biopharmaceuticals R&D, AstraZeneca, Av de Roma 81, 08029, Barcelona, Spain.

13. Protein Data Bank Japan, Institute for Protein Research, Osaka University, Suita, Osaka 565-0871, Japan.
14. Department of Biological Sciences, CW 405, Biological Sciences Building, University of Alberta, Edmonton, Alberta, Canada T6G 2E9
15. Interuniversity Institute of Bioinformatics in Brussels, VUB/ULB, Brussels, 1050, Belgium.
16. Structural Biology Brussels, AI lab, Chemistry department, Vrije Universiteit Brussel, Brussels, 1050, Belgium.
17. Wellcome Biomolecular NMR Facility, School of Natural Sciences, University of Kent, Canterbury, CT2 7NZ, United Kingdom.
18. School of Molecular Biosciences, College of Medical, Veterinary and Life Sciences, Joseph Black Building, University of Glasgow, G12 8QQ, United Kingdom.
19. Centre for Host-Microbiome Interactions, King's College London, London, SE1 9RT, UK
20. Department of Chemistry and Chemical Biology, Center for Biotechnology and Interdisciplinary Studies, Rensselaer Polytechnic Institute, Troy, NY 12180 USA.
21. Department of Chemistry and Quantum Theory Project, University of Florida, Gainesville, Florida, 32607, USA.
22. RCSB Protein Data Bank, Rutgers, The State University of New Jersey, Piscataway, NJ, USA.
23. Gdańsk University of Technology, Faculty of Electronics, Telecommunications and Informatics, G. Narutowicza 11/12, 80-233 Gdańsk, Poland
24. Faculty of Chemistry, University of Gdańsk, Wita Stwosza 63, 80-308 Gdańsk, Poland
25. Structural Biology Initiative, Advanced Science Research Center at the CUNY Graduate Center, New York, NY, USA
26. University Grenoble Alpes, CNRS, CEA, IBS, F-38000, Grenoble, France
27. Department of Chemistry & Chemical Biology, Rutgers University, Piscataway, NJ 08854, USA.
28. Molecular Biophysics and Integrated Bioimaging Division, Lawrence Berkeley National Laboratory, Berkeley, CA, USA

```
save_nef_nmr_spectrum_nnoe
   _nef_nmr_spectrum.sf_category                  nef_nmr_spectrum
   _nef_nmr_spectrum.sf_framecode                 nef_nmr_spectrum_nnoe
   _nef_nmr_spectrum.num_dimensions               3
   _nef_nmr_spectrum.chemical_shift_list          nef_chemical_shift_list_myList

   loop_
      _nef_spectrum_dimension.dimension_id
      _nef_spectrum_dimension.axis_unit
      _nef_spectrum_dimension.axis_code

      1  ppm  1H
      2  ppm  1H
      3  ppm  15N
   stop_

   loop_
      _nef_spectrum_dimension_transfer.dimension_1
      _nef_spectrum_dimension_transfer.dimension_2
      _nef_spectrum_dimension_transfer.transfer_type

      1  2  through-space
      1  3  onebond
   stop_

   loop_
      _nef_peak.index
      _nef_peak.peak_id
      _nef_peak.volume
      _nef_peak.volume_uncertainty
      _nef_peak.height
      _nef_peak.height_uncertainty
      _nef_peak.position_1
      _nef_peak.position_uncertainty_1
      _nef_peak.position_2
      _nef_peak.position_uncertainty_2
      _nef_peak.position_3
      _nef_peak.position_uncertainty_3
      _nef_peak.chain_code_1
      _nef_peak.sequence_code_1
      _nef_peak.residue_name_1
      _nef_peak.atom_name_1
      _nef_peak.chain_code_2
      _nef_peak.sequence_code_2
      _nef_peak.residue_name_2
      _nef_peak.atom_name_2
      _nef_peak.chain_code_3
      _nef_peak.sequence_code_3
      _nef_peak.residue_name_3
      _nef_peak.atom_name_3

      1    1    2.244  .    35743  .  7.375  .  7.271   .  119.59  .  .  .  .  .  .  .  .  .  .  .  .  .
      2    2    4.624  .  3669816  .  7.296  .  7.284   .  119.54  .  .  .  .  .  .  .  .  .  .  .  .  .
      3    3    1.341  .    29283  .  8.743  .  8.720   .  113.46  .  .  .  .  .  .  .  .  .  .  .  .  .
   stop_
save_
```

Metadata

Tabular data

Tabular data

Tabular data

Saveframe

**Figure S1. Overview of a `nef_nmr_spectrum` saveframe** as spectrum-specific peak lists. The example illustrates three loops containing tabular data: `nef_spectrum_dimension` (top), `nef_spectrum_dimension_transfer` (middle), and `nef_peak` list (bottom). Metadata blocks (yellow) define general information such as saveframe category and number of dimensions.

```
data_aria2_run1_it8
loop_
_atom_site.group_PDB
_atom_site.id
_atom_site.type_symbol
_atom_site.label_atom_id
_atom_site.label_alt_id
_atom_site.label_comp_id
_atom_site.label_asym_id
_atom_site.label_entity_id
_atom_site.label_seq_id
_atom_site.pdbx_PDB_ins_code
_atom_site.Cartn_x
_atom_site.Cartn_y
_atom_site.Cartn_z
_atom_site.occupancy
_atom_site.B_iso_or_equiv
_atom_site.pdbx_formal_charge
_atom_site.auth_seq_id
_atom_site.auth_comp_id
_atom_site.auth_asym_id
_atom_site.auth_atom_id
_atom_site.pdbx_PDB_model_num
_atom_site.pdbx_atom_ambiguity
```

*Sequence, residue & chain mapping*

*Model number*

*NEF atom mapping*

```
...
ATOM 1290     H HG2  . PRO A 1  94 ?  -1.680   14.783   -0.441   1.00   0.00 ? 94 PRO A HG2  1 HGx
ATOM 1291     H HG3  . PRO A 1  94 ?  -2.104   13.132    0.063   1.00   0.00 ? 94 PRO A HG3  1 HGy

ATOM 3584     H HG2  . PRO A 1  94 ?  -1.739   15.019   -0.236   1.00   0.00 ? 94 PRO A HG2  2 HGx
ATOM 3585     H HG3  . PRO A 1  94 ?  -2.037   13.312    0.161   1.00   0.00 ? 94 PRO A HG3  2 HGy

ATOM 8172     H HG2  . PRO A 1  94 ?   0.058   15.692   -0.055   1.00   0.00 ? 94 PRO A HG2  4 HGy
ATOM 8173     H HG3  . PRO A 1  94 ?  -1.697   15.427   -0.136   1.00   0.00 ? 94 PRO A HG3  4 HGx

ATOM 10466    H HG2  . PRO A 1  94 ?   0.074   15.596   -0.390   1.00   0.00 ? 94 PRO A HG2  5 HGy
ATOM 10467    H HG3  . PRO A 1  94 ?  -1.679   15.309   -0.346   1.00   0.00 ? 94 PRO A HG3  5 HGx


ATOM 991      H HB3  . SER A 1  71 ?   4.517  -14.442    9.910   1.00   0.00 ? 71 SER A HB3  1 HBy
ATOM 992      H HB2  . SER A 1  71 ?   3.441  -15.170    8.718   1.00   0.00 ? 71 SER A HB2  1 HBx

ATOM 3285     H HB3  . SER A 1  71 ?   4.349  -14.667    8.317   1.00   0.00 ? 71 SER A HB3  2 HBy
ATOM 3286     H HB2  . SER A 1  71 ?   3.917  -14.893   10.012   1.00   0.00 ? 71 SER A HB2  2 HBx

ATOM 5579     H HB3  . SER A 1  71 ?   3.853  -14.812   10.657   1.00   0.00 ? 71 SER A HB3  3 HBx
ATOM 5580     H HB2  . SER A 1  71 ?   5.247  -13.963    9.990   1.00   0.00 ? 71 SER A HB2  3 HBy

ATOM 7873     H HB3  . SER A 1  71 ?   5.000  -14.221   10.549   1.00   0.00 ? 71 SER A HB3  4 HBy
ATOM 7874     H HB2  . SER A 1  71 ?   4.011  -15.115    9.395   1.00   0.00 ? 71 SER A HB2  4 HBx
```

**Figure S2. Example of mmCIF IUPAC to NEF atom name mapping.** mmCIF tags (`_atom_site.auth_seq_id`, `_atom_site.auth_comp_id`, `_atom_site.auth_asym_id` and `_atom_site.pdbx_atom_ambiguity`) map atom coordinates with NEF atom names.

```
save_nef_nmr_meta_data
   _nef_nmr_meta_data.sf_category              nef_nmr_meta_data
   _nef_nmr_meta_data.sf_framecode             nef_nmr_meta_data
   _nef_nmr_meta_data.format_name              nmr_exchange_format
   _nef_nmr_meta_data.format_version           1.1
   _nef_nmr_meta_data.program_name             ARIA
   _nef_nmr_meta_data.program_version          2.3.3
   _nef_nmr_meta_data.creation_date            2023-12-07T11:11:40.108474
   _nef_nmr_meta_data.uuid                     ARIA-2023-12-07T11:11:40.108474-721040
   _nef_nmr_meta_data.coordinate_file_name     .
   _nef_nmr_meta_data.aria_project_name        ntd
   _nef_nmr_meta_data.aria_run                 run1
   _nef_nmr_meta_data.aria_iteration           8
```

***Additional name-space tag-value pair***

```
   loop_
      _nef_program_script.program_name
      _nef_program_script.script_name

      ARIA NEFio.py

   stop_

   loop_
      _nef_run_history.run_number
      _nef_run_history.program_name
      _nef_run_history.program_version
      _nef_run_history.aria_input_uuid
```

***Additional name-space loop tag***

```
      1    AnalysisStructure 3.2.1 AnalysisStructure-2023-12-06T13:22:52.020106-1222356005

   stop_

save_
```

***Additional name-space saveframe***

```
save_aria_violation_list
   _aria_violation_list.sf_category      aria_violation_list
   _aria_violation_list.sf_framecode     aria_violation_list

   loop_
      _aria_violation.index
      _aria_violation.aria_id
      _aria_violation.chain_code_1
      _aria_violation.sequence_code_1
      _aria_violation.residue_name_1
      _aria_violation.atom_name_1
      _aria_violation.chain_code_2
      _aria_violation.sequence_code_2
      _aria_violation.residue_name_2
      _aria_violation.atom_name_2
      _aria_violation.weight
      _aria_violation.target_value
      _aria_violation.lower_limit
      _aria_violation.upper_limit
      _aria_violation.calc_dist
      _aria_violation.calc_dist_error
      _aria_violation.lower_bound_violation
      _aria_violation.upper_bound_violation
      _aria_violation.frac_viol
      _aria_violation.used_for_calculation
      _aria_violation.nef_restraint_id
      _aria_violation.nef_list_name

      1          0 A    5    THR H    A    4    TYR HA   1.000  2.161  1.577  2.745  3.105  0.423  0.000  0.577  0.71  true  1      nef_distance_restraint_list_1
      2         17 A    3    ILE H    A    18   PRO HA   1.000  2.084  1.541  2.626  3.018  0.499  0.000  0.391  0.86  true  18     nef_distance_restraint_list_1
      3         25 A    25   THR H    A    15   THR HB   1.000  3.598  1.980  5.216  7.666  1.855  0.000  2.879  0.86  true  26     nef_distance_restraint_list_1
      4         25 A    27   LEU H    A    15   THR HB   1.000  3.598  1.980  5.216  7.666  1.855  0.000  2.879  0.86  true  26     nef_distance_restraint_list_1
```

**Figure S3. Example of NEF namespace-specific data.** NEF supports extra namespace-specific metadata tags, additional loop columns, new loops, or even entirely new saveframes. In the example shown, Aria uses the '`_aria`' label for additional data such as the name-space tag-value pair `_nef_nmr_meta_data.aria_project_name` and the additional namespace saveframe `save_aria_violation_list`.

2K3A – data-rich protein

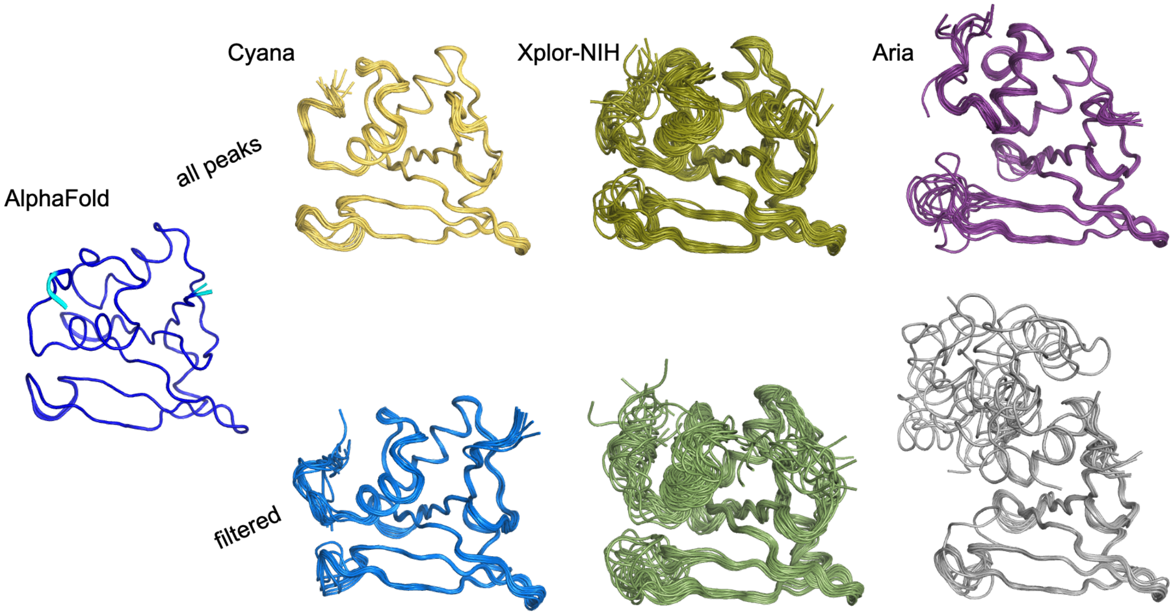


**Figure S4. Structural ensembles calculated for case-1 (2K3A).** Top row shows the ensembles generated by Cyana, Xplor-NIH and Aria, respectively, using the unfiltered peak dataset. Bottom row shows the ensembles generated using the filtered peak dataset. The AlphaFold3 predicted structure (*68*) coloured by plDDT score (blue: plDDT > 90, cyan: 90 > plDDT > 70, yellow: 70 > plDDT > 50, orange: plDDT < 50) has been included for reference. Only residues 47-155 of all ensembles are shown for clarity.

All models in each of the ensembles are superimposed on the first model using the backbone heavy atoms (O, N, CA, C) and the secondary structure regions (alpha helices: 57-64, 71-73, 76-86, 143-146; beta strands: 89-91, 98-101, 110-116, 122-127, 136-141, 150-153) as determined for the AlphaFold3 model by the DSSP algorithm (*69*) implemented in CcpNmr AnalysisStructure.

2M47 – limited-data protein

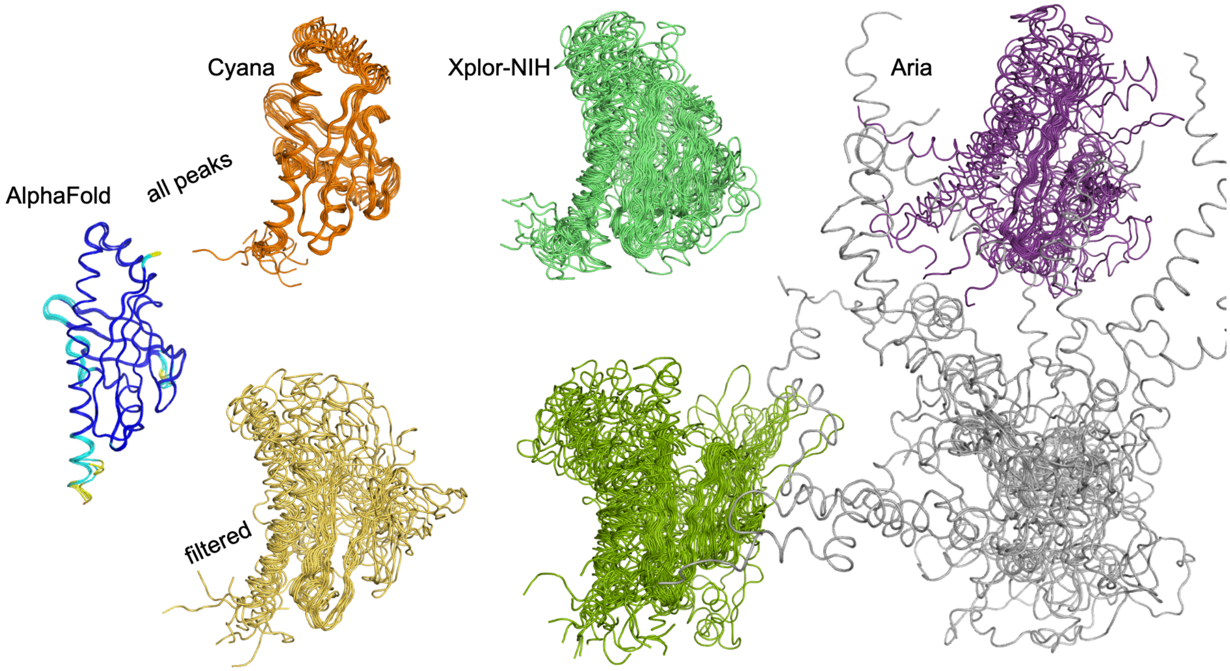


**Figure S5. Structural ensembles calculated for case-2 (2M47).** Top row shows the ensembles generated by Cyana, Xplor-NIH and Aria, respectively, using the unfiltered peak dataset. Bottom row shows the ensembles generated using the filtered peak dataset. The AlphaFold3 predicted structure (*68*) coloured by plDDT score (blue: plDDT > 90, cyan: 90 > plDDT > 70, yellow: 70 > plDDT > 50, orange: plDDT < 50) has been included for reference.

All models in each of the ensembles are superimposed on the first model using the backbone heavy atoms (O, N, CA, C) and, due to the large structural disorder, the alpha-helical regions only (alpha helices: 5-13, 79-84, 89-98, 101-110) as determined for the AlphaFold3 model by the DSSP algorithm (*69*) implemented in CcpNmr AnalysisStructure.

short peptide – underdetermined

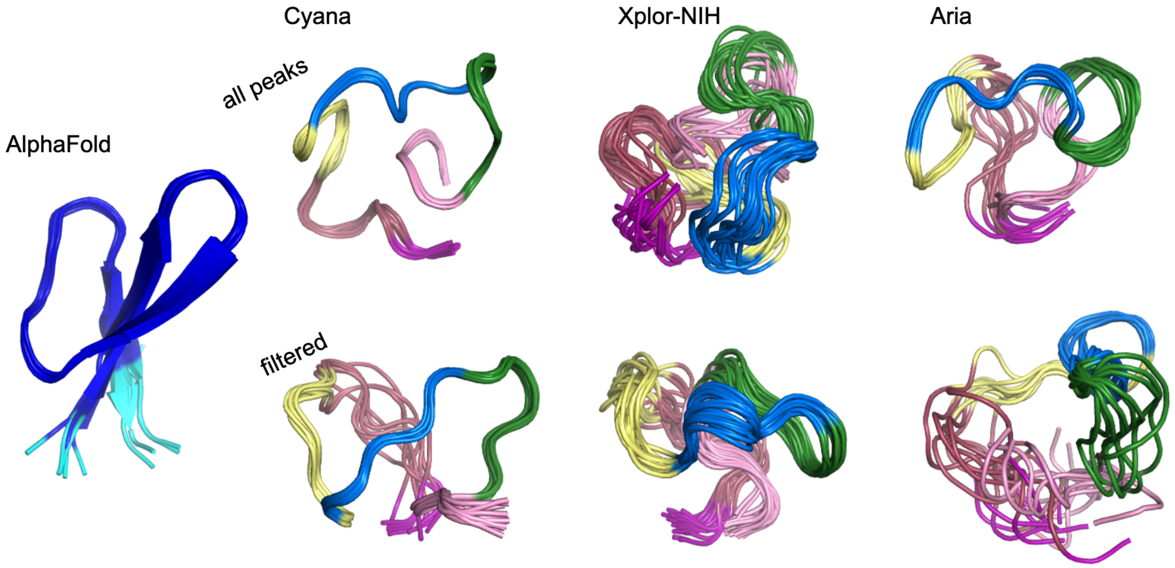


**Figure S6. Structural ensembles calculated for case-3 (peptide).** Top row shows the ensembles generated by Cyana, Xplor-NIH and Aria, respectively, using the unfiltered peak dataset. Bottom row shows the ensembles generated using the filtered peak dataset. The AlphaFold3 predicted structure (*68*) coloured by plDDT score (blue: plDDT > 90, cyan: 90 > plDDT > 70, yellow: 70 > plDDT > 50, orange: plDDT < 50) has been included for reference. The calculated Cyana, Xplor-NIH and Aria ensembles have been coloured by sequence to facilitate comparison.

All models in each of the ensembles are superimposed on the first model using the backbone heavy atoms (O, N, CA, C) and using all residues.

**Format specific details**

The universally unique identifier (UUID) has the form `<program-name>-<time-stamp>-<random-integer>`, with `<program-name>` denoting the last program to alter the file, `<time-stamp>` recommended to be ISO 8601 with microsecond precision, e.g. 2016-07-20T18:25:26.324290, and `<random-integer>` to have 10 digits.

Data items that include spaces, tabs, or newlines must be enclosed in single or double quotes, or specified by a multi line quote that starts and ends with a `<semi-colon>` (;) as the first character of a line. For multi-line strings delimited by a `<new-line><semi-colon>`, or that start with a `<semi-colon>` it is conventional to consistently space indent the complete line to avoid early string termination.

Atoms can either be added or omitted in the `_nef_sequence` loop using +`<atom_name>` or –`<atom_name>` identifiers.

Namespaces are to be used with caution and to follow common sense. NEF-defined saveframes and loops can be augmented as illustrated by the examples. Likewise, the NEF specification currently does not prohibit augmenting saveframes and loops defined by other namespaces, e.g. aria or meld, in a similar fashion. However, it is highly recommended not to do this, as confusion is likely to arise. Whereas NEF tags are defined by the NEF dictionary and thus can be probed for, this is not a requirement for tags added through the namespace mechanism. A clarification and tightening of the namespace mechanism will be initiated.

**Table ST2. NEF testing data sets**

| PDB ID | BMRB ID | NESG ID | Number of restraint saveframes | Sequence length | Comments | PDB Deposition Authors |
|---|---|---|---|---|---|---|
| 1PQX | 5844 | ZR18 | 2 | 87 | | Baran, M.C., Aramini, J.M., Xiao, R., Huang, Y.J., Acton, T.B., Shih, L., Montelione, G.T. |
| 2JR2 | 15317 | CsR4 | 2 | 2 x 76 | dimer | Ramelot, T.A., Cort, J.R., Wang, H., Nwosu, C., Cunningham, K., Owens, L., Ma, L.-C., Xiao, R., Liu, J., Baran, M.C., Swapna, G., Acton, T.B., Rost, B., Montelione, G.T., Kennedy, M.A. |
| 2JUW | 15456 | SoR77 | 2 | 2 x 80 | dimer | Ramelot, T.A., Cort, J.R., Wang, D., Nwosu, C., Owens, L., Xiao, R., Liu, J., Baran, M.C., Swapna, G.V.T., Acton, T.B., Rost, B., Montelione, G.T., Kennedy, M.A. |
| 2K2E | 157021 | BeR31 | 2 | 158 | | Cort, J.R., Ho, C.K., Nwosu, C., Maglaqui, M., Xiao, R., Liu, J., Baran, M.C., Swapna, G., Acton, T.B., Rost, B., Montelione, G.T., Kennedy, M.A. |
| 2KCU | 16097 | CtR107 | 2 | 166 | | Mills, J.L., Zhang, Q., Sukumaran, D.K., Wang, D., Jiang, M., Foote, E.L., Xiao, R., Nair, R., Everett, J.K., Swapna, G.V.T., Acton, T.B., Rost, B., Montelione, G.T., Szyperski, T. |
| 2KKO | 16368 | MbR242E | 2 | 2 x 108 | dimer | Ramelot, T.A., Cort, J.R., Wang, D., Ciccosanti, C., Jiang, M., Nair, R., Rost, B., Swapna, G., Acton, T.B., Xiao, R., Everett, J.K., Montelione, G.T., Kennedy, M.A. |
| 2KO1 | 16486 | CtR148A | 2 | 2 x 83 | dimer | Eletsky, A., Garcia, E., Wang, H., Ciccosanti, C., Jiang, M., Nair, R., Rost, B., Acton, T.B., Xiao, R., Everett, J.K., Lee, H., Prestegard, J.H., Montelione, G.T., Szyperski, T. |
| 2KO7 | 16406 | NA | 2 | 175 | | Zheng, S., Leeper, T., Varani, G., |
| 2KPU | 16570 | DhR29B | 2 | 96 | | Cort, J.R., Ramelot, T.A., Yang, Y., Belote, R.L., Ciccosanti, C., Haleema, J., Acton, T.B., Xiao, R., Everett, J.K., Montelione, G.T., Kennedy, M.A. |
| 2KW5 | 16806 | SgR145 | 4 | 202 | RDC restraints | Rossi, P., Forouhar, F., Lee, H., Lange, O., Mao, B., Lemak, A., Maglaqui, M., Belote, R., Ciccosanti, C., Foote, E., Sahdev, S., Acton, T., Xiao, R., Everett, J., Baker, D., Montelione, G.T. |

| | | | | | | |
|---|---|---|---|---|---|---|
| 2KZN | 17008 | SR10 | 3 | 147 | RDC restraints | Ertekin, A., Maglaqui, M., Janjua, H., Cooper, B., Ciccosanti, C., Rost, B., Acton, T.B., Xiao, R., Everett, J.K., Prestegard, J., Lee, H., Aramini, J.M., Rossi, P., Montelione, G.T. |
| 2LOY | 16833 | WR73 | 3 | 183 | RDC restraints | Aramini, J.M., Rossi, P., Cort, J.R., Lee, H., Janjua, H., Maglaqui, M., Cooper, B., Xiao, R., Acton, T.B., Everett, J.K., Montelione, G.T. |
| 2LUZ | 18547 | MiR12 | 2 | 182 | | Ramelot, T.A., Yang, Y., Lee, H., Pederson, K., Lee, D., Kohan, E., Janjua, H., Xiao, R., Acton, T.B., Everett, J.K., Wrobel, R.L., Bingman, C.A., Singh, S., Thorson, J.S., Prestegard, J.H., Montelione, G.T., Phillips Jr., G.N., Kennedy, M.A. |
| 2PNG | 15449 | NA | 1 | 89 | no dihedral angle restraints | Płoskoń, E.A., Arthur, C.J., Evans, S.E., Williams, C., Crosby, J., Crump, M.P. |
| 6NBN | 30550 | NA | 2 | 123 | Protein-ligand complex | Jones, D.N., Wang, J. |
